# Ultra-rapid metagenotyping of the human gut microbiome

**DOI:** 10.1101/2020.06.12.149336

**Authors:** Zhou Jason Shi, Boris Dimitrov, Chunyu Zhao, Stephen Nayfach, Katherine S. Pollard

**Affiliations:** Chan Zuckerberg Biohub, Data Science, San Francisco, CA; Gladstone Institutes, San Francisco, CA; Chan Zuckerberg Initiative, CA; Department of Energy, Joint Genome Institute, Walnut Creek, CA; Lawrence Berkeley National Laboratory, Environmental Genomics and Systems Biology Division, Berkeley, CA; University of California San Francisco

## Abstract

Sequence variation is used to quantify population structure and identify genetic determinants of phenotypes that vary within species. In the human microbiome and other environments, single nucleotide polymorphisms (SNPs) are frequently detected by aligning metagenomic sequencing reads to catalogs of genes or genomes. But this requires high-performance computing and enough read coverage to distinguish SNPs from sequencing errors. We solved these problems by developing the GenoTyper for Prokaytotes (GT-Pro), a suite of novel methods to catalog SNPs from genomes and use exact k-mer matches to perform ultra-fast reference-based SNP calling from metagenomes. Compared to read alignment, GT-Pro is more accurate and two orders of magnitude faster. We discovered 104 million SNPs in 909 human gut species, characterized their global population structure, and tracked pathogenic strains. GT-Pro democratizes strain-level microbiome analysis by making it possible to genotype hundreds of metagenomes on a personal computer.

**Software availability:** GT-Pro is available at https://github.com/zjshi/gt-pro.

## Introduction

Microbial species harbor extensive genetic variation, including single nucleotide polymorphisms (SNPs), structural variants (SVs), and mobile genetic elements. SNPs in particular are useful for population genetic analyses1, such as tracking transmission of strains between environments or locations, reconstructing strain phylogenetic relationships, resolving mixtures of genotypes within a host, and depicting population diversity or structure along environmental gradients. Additionally, SNPs can result in changes in protein function. For example, a single SNP in the Dadh gene of the human commensal *Eggerthella lenta* can predict activity of levodopa, the primary medication used to treat Parkinson’s disease^2^. Quantifying intra-species genomic variation in the human microbiome is a prerequisite to the potential application of microbiome genomics to precision medicine.

Several approaches exist for identifying SNPs in microbiomes. The gold standard^3^ is to sequence individual isolate genomes and identify mismatches in whole-genome alignments. In contrast, metagenomes are a rich source of strain level diversity for uncultivated taxa. In a landmark study, Schloissnig et al^4^. discovered 10.3 million SNPs for 101 human gut species by aligning short reads from shotgun metagenomes to reference genomes. This approach is known as “metagenotyping” and has since been featured in several tools, including Constrains^5^, MIDAS^6^, metaSNV^7^, DESMAN^8^ and StrainPhlAn^9^. While algorithms for read alignment have improved, the approach is still computationally costly. Exact matching algorithms such as Kraken^10^, CLARK^11^ and bfMEM^12^, have been developed as a more efficient solution to the read mapping problem, achieving speedups by orders of magnitude. However, these tools have thus far been used to quantify the abundance of microbiome taxa, rather than identify intra-species genetic variation. Genotyping by exact matches between reads and short sequences covering SNPs was implemented in the method LAVA^13^ for human whole-genome sequencing data. Our goal was to extend this approach to metagenomes by addressing the challenges presented by complex mixtures of species and strains within microbiome samples, while also making software that could run on a personal computer.

## Results

### A novel framework for *in silico* genotyping of microbiome species

We introduce the GenoTyper for PROkaryotes (GT-Pro), which is a novel computational pipeline that utilizes an exact matching algorithm to perform ultra-rapid and accurate genotyping of known SNPs from metagenomes. Our proof-of-principle initial implementation of this approach focuses on the human gut microbiome. We created a reference database of 104 million common SNPs that we identified using 112,904 high-quality genomes from 909 human gut microbiome species. Then we used this catalog to perform reference-based SNP calling for 25,133 publicly available metagenomes, providing insight into strain variation across individuals and geographic regions. Our results demonstrate the feasibility of performing large-scale metagenotyping without need for high-performance computing.

To overcome the low throughput, sensitivity and species coverage of current alignment-based metagenotyping methods, we developed the GT-Pro framework (Fig. 1). Our key innovations are (i) capturing the majority of common variation found in microbiome genomes with a compact database of SNP covering k-mers (sck-mers), (ii) selecting highly species-specific sck-mers, overcoming high false positives associated with k-mer exact-matching methods, and (iii) developing and optimizing algorithms and data structures for exact matching of metagenomic sequencing reads to these sck-mers, enabling SNPs to be detected rapidly and accurately in microbiome samples. Building a version of GT-Pro for a given environment involves 1) discovering common SNPs in assembled genomes for each species, 2) optionally identifying linkage disequilibrium (LD) blocks and “tag” SNPs that capture most variation within each block, and 3) designing species-specific sck-mers. We focus on common SNPs) because this allows us to create a virtual genotyping “array” that is a data structure small enough to fit in computer memory while stll capturing the majority of prevalent genetic variation for each of many species.

**Figure 1.**
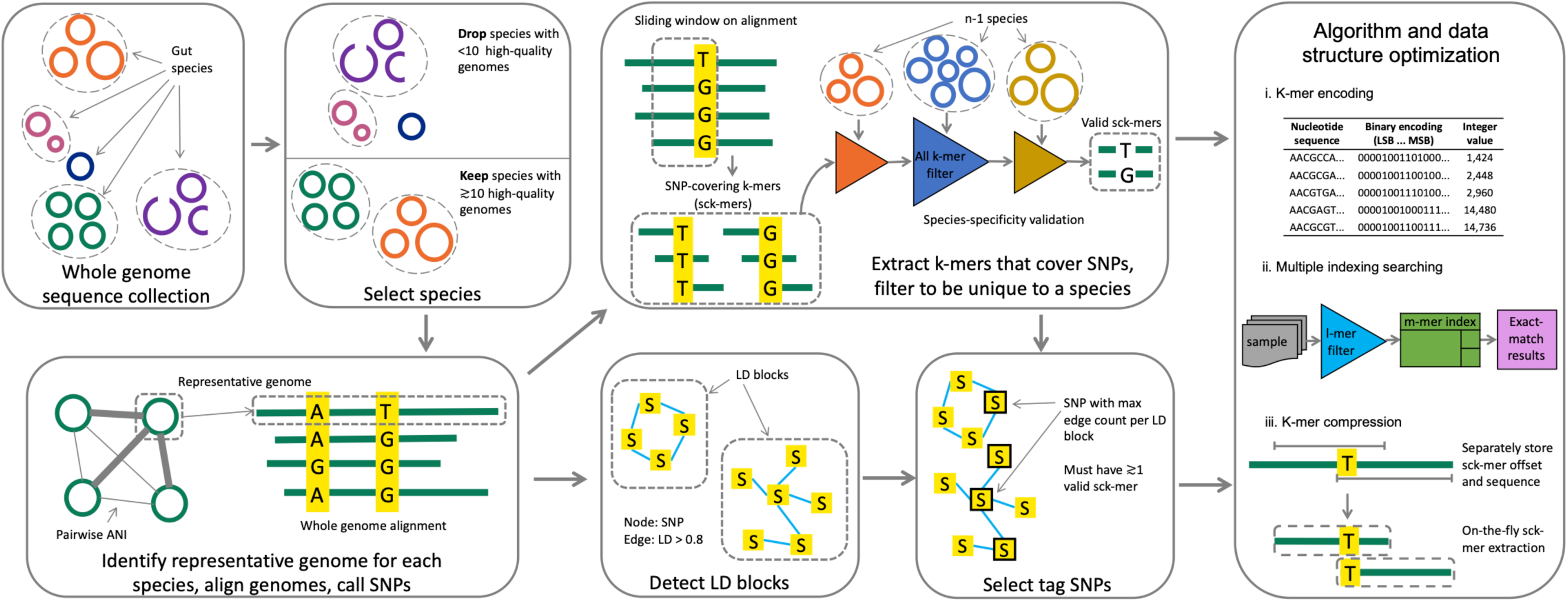
In sillico metagenotyping framework. Our method starts with a whole genome sequence collection and identifies species with sufficient high-quality genomes to call SNPs. For each species, a representative genome is chosen based on pairwise Average Nucleotide Identity (ANI) plus assembly quality metrics. SNPs are called per species based upon whole genome alignment of conspecific genomes to the representative genome. Common (MAF > 1%) bi-allelic SNPs are selected for genotyping. Up to 4 X k candidate k-mers are extracted per SNP site, covering both the reference and alternative allele on forward and reverse complementary strands (sck-mers; k=31 in this study). These candidate sck-mers are iteratively filtered through species-specificity filters of all unique k-mers present in the genomes of every other species, including species with insufficient high-quality genomes for genotyping. Only SNPs with sck-mers for both the reference and alternative allele are retained. SNPs are clustered based on pairwise linkage disequilibrium (LD). LD blocks are detected with a threshold of mean r2 > 0.81, and we select a tag SNP with species-specific sck-mers and the highest LD to other SNPs in the block. Optimized algorithms and compressed representations of sck-mer data enable rapid metagenotyping. Further details in Methods and Figure 3.

### A database of common SNPs for bacterial species in the gut microbiome

As a case study, we applied GT-Pro to the human gut microbiome due to the large number of microbial genomes from this environment and its important role in human health. To construct a SNP catalog, we used 112,904 high-quality genomes (>= 90% completeness and <= 5% contamination^14^) from 909 species (minimum = 10 genomes, median = 35 genomes) that we downloaded from the Unified Gastrointestinal Genomes (UHGG) resource^15^ (Fig. S1, S2 and Table S1). These include both metagenome-assembled genomes^16–18^ (i.e. MAGs, 94.1%) as well as cultivated isolates (5.9%) and were derived from geographically and phenotypically diverse human subjects. We performed whole-genome alignments for each species, revealing 104,171,172 common, core-genome SNPs (minor allele frequency >= 1%, site prevalence >= 90%), the vast majority of which (93.4%) were bi-allelic (Fig. 2a, S3a and S4). An extremely low fraction of SNPs (<0.2%) either disrupted a stop codon or introduced a premature one, which is one indicator of false positives (Fig. 2a). For context, this catalog is 10-fold larger than the one established by Schloissnig et al. and 1.22-fold larger than the catalogue of all human SNPs^19^ (Fig. S1). Consistent with previous reports^4^, SNP density, nucleotide diversity, and the rate of nonsynonymous versus synonymous mutations (pN/pS) varied across species and phyla (Fig. 2b and Fig. S5-8), which may reflect differences in selective pressures, population sizes, or transmission modes.

**Figure 2.**
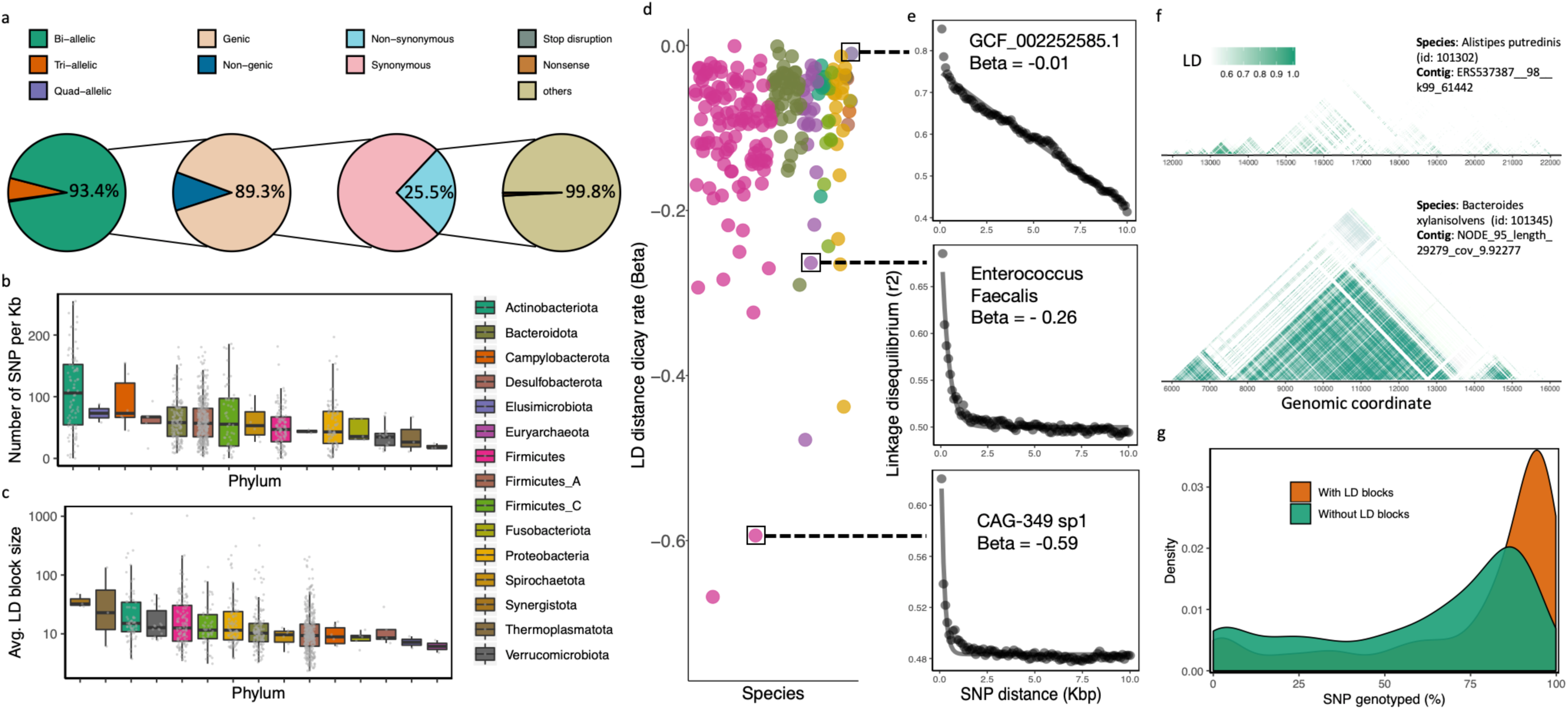
Genetic landscape of 909 human gut species. (a) Summary of common SNP characteristics across all species (from left to right): at most SNPs only two alleles are observed, bi-allelic SNPs are mostly within protein-coding genes, these are largely synonymous, and the non-synonymous ones rarely disrupt or introduce a stop codon. (b and c) Phyla differ in their median SNP density and average LD block size with significant variation in density across species within each phylum. (d) Rate of LD distance decay across gut bacterial species. (b-e) are colored by bacterial phylum and share the same color scheme. (e) Examples of LD distance decay for individual species. From top to bottom are three species (species id: 102446, 101694 and 102831) with increasingly fast LD distance decay, suggesting higher recombination rates. Curves represent the fitted exponential decay model. (f) Visualization of two distinct haplotype landscapes from (upper) species *Alistipes putredinis* (species id: 101302) and (lower) *Bacteroides xylanisolvens* (species id: 101345). Base axis represents and is ordered by genomic coordinate. Color indicates magnitude of LD between pairs of SNPs. The examples have the same genomic span (10,000 bp). (g) Distribution across species of the percentage of SNPs that can be genotyped by GT-Pro either directly (“without LD blocks”) or by imputation using genotyped tag SNPs (“with LD blocks”). For a typical species, ∼75% of SNPs can be genotyped directly and ∼95% can be imputed.

We hypothesized that the SNP database could be greatly compressed by clustering SNPs into linkage disequilibrium (LD) blocks that co-vary across reference genomes (Fig. S9) and selecting a single “tag” SNP per LD block. A similar strategy is commonly used when designing genotyping arrays in human genetics. Using single-linkage clustering (R^2^ > 0.81), the 104 million SNPs were clustered into 6.8 million LD blocks, representing a >15-fold reduction in database size and revealing a remarkable degree of local genomic structure. Our choice of R^2^ is motivated by thresholds used for high confidence SNP imputation in other species and the fact that discovery of LD blocks stabilizes in this range for gut species (Fig. S10). On average LD blocks spanned ∼4.3Kbp and ∼23.5 SNPs, though the number and size of LD blocks varied considerably across bacterial species (Fig. 2c, S5c and S11a and b). As expected, linkage between SNPs decayed with increasing genomic distance (Fig. 2d-f), though decay rates differed substantially across species (Fig. 2d-e). Altogether, these differences in genetic diversity and structure across species likely reflect variation in recombination rates and/or the number and relatedness of sequenced genomes.

### Species-specific kmers enable accurate and efficient identification of SNPs

Having constructed a large SNP catalog of the gut microbiome, we next used GT-Pro to identify k-mers that could unique identify each SNP from shotgun metagenomes. We empirically determined that length k=31 ensured high specificity while limiting compute and memory requirements. Of the ∼13.3 billion candidate 31-mers that overlapped a SNP (124 per SNP), we identified 5.7 billion that were unique. These kmers overlapped 51% of the 104 million SNPs for 65% of LD blocks (mean 108 sck-mers per SNP, >1 sck-mer for 97% of species, Fig. S1 and S12). We refer to these as species-specific, SNP-covering kmers (sck-mers). Species with few or no SNPs that can be genotyped with this strategy include those with a very close relative and are most common within Actinobacteria (Fig. 2g and S3b). While only 50% of SNPs were tagged by a sck-mer, they capture 83% of the within species variation compared to whole-genome average nucleotide identity, and achieve a much higher level of resolution compared to individual taxonomic markers (Fig. S13). Due to the large scale of the database, GT-Pro uses a highly efficient data structure to store the sck-mers, requiring only 13 GB of RAM and permitting GT-Pro to run on most modern personal computers (Fig. S14 and S15). We also created a low memory version of the GT-Pro database (< 4 GB RAM) which just stores sck-mers for a single “tag” SNP per LD block (Methods) and still captures the majority of within species variation (Fig. S13).

### Optimized k-mer exact matching accelerates metagenotyping 100-fold

To search for exact matches between billions of k-mers among metagenome reads and billions of sck-mers in the GT-Pro database, it is crucial to have a highly efficient search algorithm with low RAM and I/O requirements. To this end, we developed an exact match algorithm that leverages data structures optimized for this specific application (Fig. S16). Our approach is similar to a multi-index search with three main steps operating on bit encoded k-mers (2 bits per base) (Fig. S16a). After generating all k-mers in each metagenomic sequencing read, GT-Pro uses a l-bit Bloom filter on the first l<k bits of each k-mer to quickly rule out the vast majority of read k-mers that have no chance to match database sck-mers because they do not share an l-mer. For the k-mers that pass through the l-bit filter, the algorithm recruits an m-bit (last m bits of encoded k-mer) index to serve as secondary filter that locates a bucket of pre-sorted sck-mers in the database containing all possible exact matches to the full k-mer. Finally, the algorithm invokes a sequential search for exact matches between the full k-mer and these only the sck-mers in this bucket.

We next evaluated GT-Pro computational performance. First, we measured both speed and peak RAM use while tuning the values of l and m, two parameters derived from the l-bit and m-bit filter that are expected to have a large impact on performance due to their direct relationships with query speed and peak RAM use. In general, both performance metrics increase with higher values of l and m (Fig. 3a). Within the range of the tested parameters, we found best speed and peak RAM use with l = 30 and m = 35 in the laptop environment (26.5GB RAM) and the with l = 32 and m = 36 on a server (56.55 GB RAM). In a boundary case (l = 30 and m = 36) on the laptop where the peak RAM use hit the hardware limit, speed drops >87%. These results demonstrate that the values of l and m should be carefully chosen based on the hardware for optimal performance, which is handled automatically by GT-Pro.

**Figure 3.**
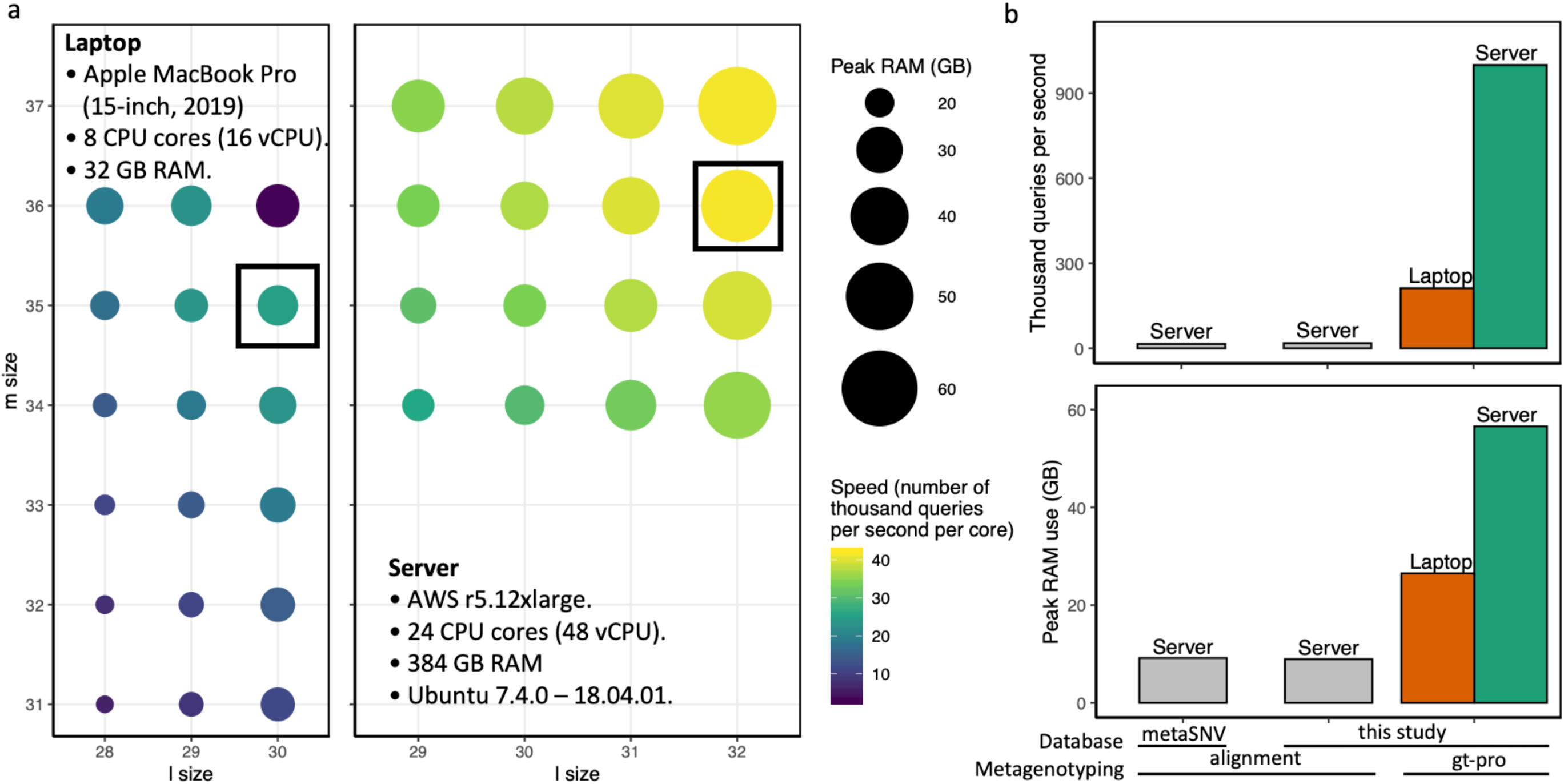
Computational performance evaluation of GT-Pro. (a) Computational performance of GT-Pro in laptop (left) and server (right) environments across values for l (Bloom filter size parameter) and m (m-index size parameter). Color gradient: processing speed, circle size: peak RAM use, black box: optimal l and m for each computing environment. (b) Comparison of speed (upper) and peak RAM usage (lower) between GT-Pro and alignment-based metagenotyping (metaSNV and MIDAS; see methods). We ran GT-Pro on both server (green) and laptop (yellow) environments, while alignment-based methods were run only in the server (grey) environment due to not being optimized for personal computers. Peak RAM usage exceeds RAM needed to store the database due to intermediate calculations, such as applying filters.

We then compared the computational performance of GT-Pro to traditional read alignment method as baseline (Fig. 3b). We arbitrarily selected a total of 40 stool metagenomes from a Tanzanian cohort^20^ (Table S7) for the evaluation. For alignment, SNPs were called mapping reads to database from GT-Pro and an independent one (metaSNV^7^). Although GT-pro had a larger peak RAM use than alignment method (<10GB), was 100x faster on a server and 10x faster on a laptop where peak RAM use was 26.5GB (Fig. 3b).

### Accurate identification of SNPs from simulated metagenomes

We next evaluated the accuracy of SNP calling with GT-Pro compared to alignment using simulated metagenomes. Towards this goal, we generated Illumina sequencing reads *in silico* from 978 human gut isolates^21^ and identified the ground truth set of SNPs based on whole-genome alignment. We first simulated reads from individual isolates with sequencing coverages ranging from 0.001x to 15x (Table S2). In this simplified scenario, genotyping errors can result from sequencing errors, insufficient coverage, or incorrect read or k-mer mapping. Across isolates and coverage levels, the false discovery rate (FDR) of genotypes was on average lower for GT-Pro (median= 0.7%, IQR=1.1%) compared to read alignment (median= 2.2%, IQR=4.7%) (Fig. 4a) while the median sensitivity of GT-Pro tended to be consistently higher (4.1-17.6%) at all coverages (Fig. 4b). While read alignment methods typically use a minimum coverage threshold to avoid false positives from sequencing error (e.g. >10x), that would have further decreased the sensitivity in this experiment.

**Figure 4.**
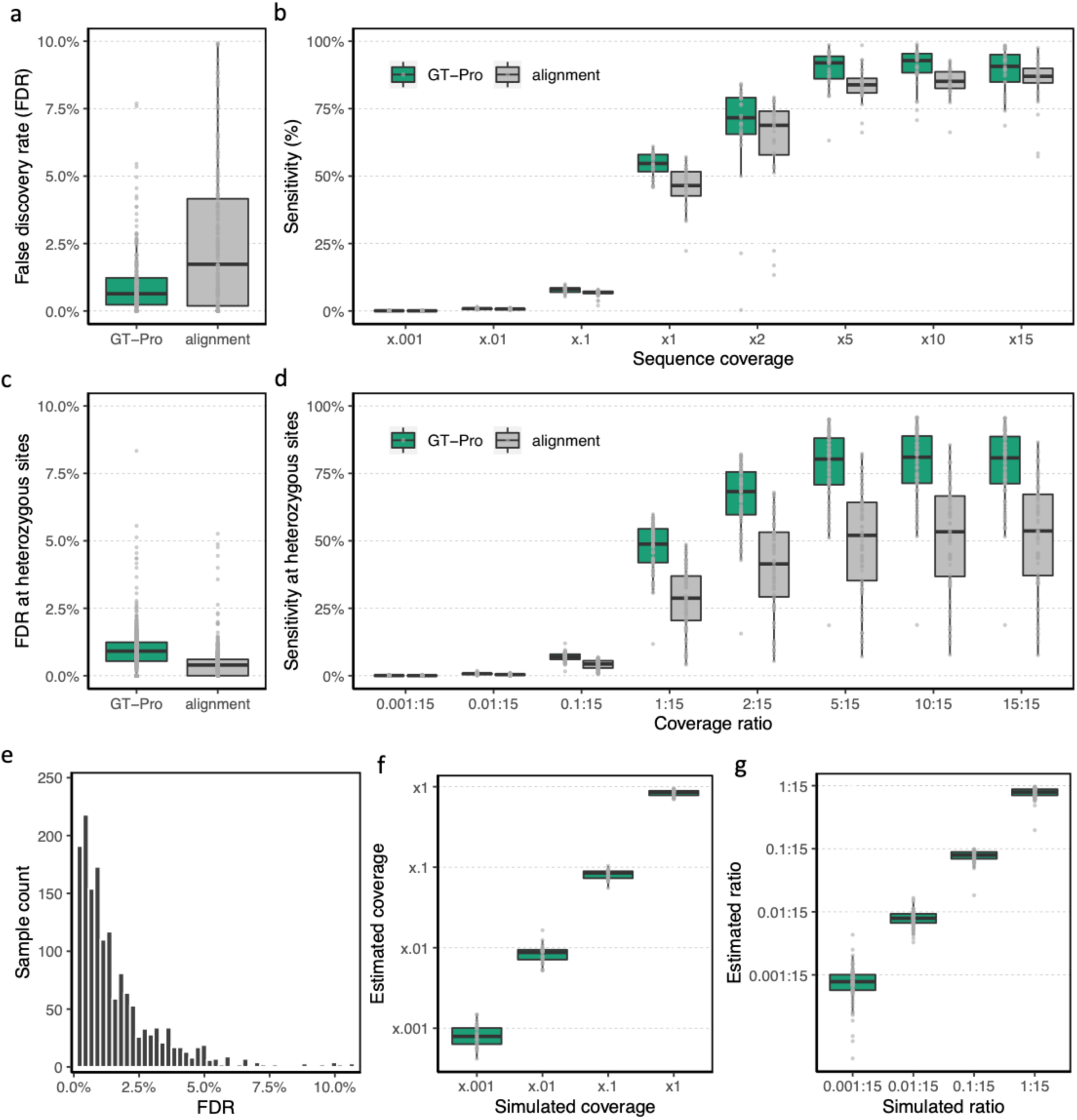
Metagenotyping accuracy evaluation of GT-Pro using simulations. Accuracy comparisons of GT-Pro and alignment-based metagenotyping across species based on reads simulated from isolate genomes with sequencing error. (a) False discovery rate at a combination of sequencing coverage ranged from 0.001x to 15x. Each observation is the result from a metagenome containing reads from one isolate. False discoveries are genotype calls that do not match the genome from which reads were simulated. (b) Sensitivity across coverage levels from the simulations in (a). Sensitivity is the probability of detecting SNPs present in the isolate genome. (c) False discovery rate at heterozygous sites in metagenomes containing reads from two isolates of each species. A combination of sequencing coverage ratio between two isolates was simulated by fixing a more abundant isolate at 15x coverage in all simulations, and varying the other isolate’s coverage from 0.001x to 15x (coverage ratio = 0.001:15 to 15:15). (d) Sensitivity at heterozygous sites in metagenomes from (c). Sensitivity is the probability of correctly calling the heterozygous genotype of sites that differ between the genomes from which reads were simulated. (e) False discovery rate of genotypes imputed from tag SNPs based on allele matching in simulations in (a). Imputation is simply done by selecting the genotype associated with the observed tag SNP. (f) Sequencing coverage estimated using read counts at GT-Pro genotyped SNPs correlates with the simulated coverage, even when coverage is <1x. Each observation is the estimate from metagenomic reads simulated with sequencing error from a single isolate genome. (g) Sequencing coverage ratio estimates based on read counts for each allele at GT-Pro genotyped heterozygous sites correlate with the simulated ratios of two isolate genomes, even when one is much less abundant than the other (<=1:15). The more abundant isolate is at 15x coverage in all simulations.

Next, we simulated metagenomes containing pairs of conspecific isolates to evaluate performance on samples with strain mixtures, exploring a range of coverage ratios from 0.001x to 15x, where one strain is always at 15x coverage and the other varying (Table S3 and S4). In terms of detecting heterozygous sites (strains with different alleles), the false discovery rate (FDR) of GT-Pro (median= 0.9%, IQR=0.7%) was slightly higher compared to alignment (median= 0.3%, IQR=0.6%) (Fig. 4c), however, median sensitivity was higher (50.5-81.6%) for GT-Pro at all coverage ratios (Fig. 4d). A higher FDR for GT-Pro is likely caused by sequencing errors that match the alternative allele by chance, which also could cause a slightly lower sensitivity for GT-Pro at homozygous sites (Fig. S17).

To evaluate genotype calls imputed from tag SNPs, we found low FDR<5% comparing to true genotypes for the vast majority of isolates (>95%) (Fig. 4e). SNPs belonging to an LD block were 5 times more likely to be detected (non-zero read count) when their tag SNPs were also detected than when they were not (Fig. S18). To show that GT-Pro is highly quantitative, we compared average coverage at SNPs in the GT-Pro output to the known genome coverage using metagenomes we simulated from individual isolates and pairs of conspecific isolates. Even at low sequencing coverage (<1X), GT-Pro was able to accurately estimate the true coverage of each species (Fig. 4f) and the ratio between two strains (Fig. 4g). These results suggest that GT-Pro allele calls and counts could be used to impute genotypes and estimate relative abundances of species and strains accurately.

### Accurate metagenotyping and gene imputation from gut metagenomes

To compare GT-Pro to existing approaches, we metagenotyped gut metagenomes^16,20,22,23^ (Table S5-10) with alignment and compared the number of genotyped SNPs plus estimates of allele frequencies and genetic distances. We found that GT-Pro genotyped more species and SNPs per metagenome (Fig. S19a-c), despite being limited to species with ≥10 genomes. This is likely due to GT-Pro having better sensitivity for low coverage species and using a human gut focused database (comparing to metaSNV). For species genotyped by both methods, within-sample heterozygosity (Fig. S18) and across-sample allele presence and frequency (Fig. 5a-d) were highly correlated. For high coverage species, alignment method detected some SNPs absent from the GT-Pro database, whereas GT-Pro detected more sites as polymorphic in medium and low coverage species (Fig. 5a-d). Despite these differences in genotyped sites, GT-Pro and alignment produce highly similar estimates of pairwise genetic distances (Jaccard index) between samples, likely because rare variants missed by GT-Pro but with sufficient coverage to be genotyped with alignment-based methods represent a small fraction of overall genetic diversity. For comparison, we repeated this analysis using only SNPs in the 16S gene and observed much lower genetic differences between samples (Fig. S21), emphasizing that GT-Pro provides strain resolution close to that of alignment and greatly exceeding that of marker gene approaches. Altogether these results are consistent with our simulations and underscore the high sensitivity of GT-Pro.

**Figure 5.**
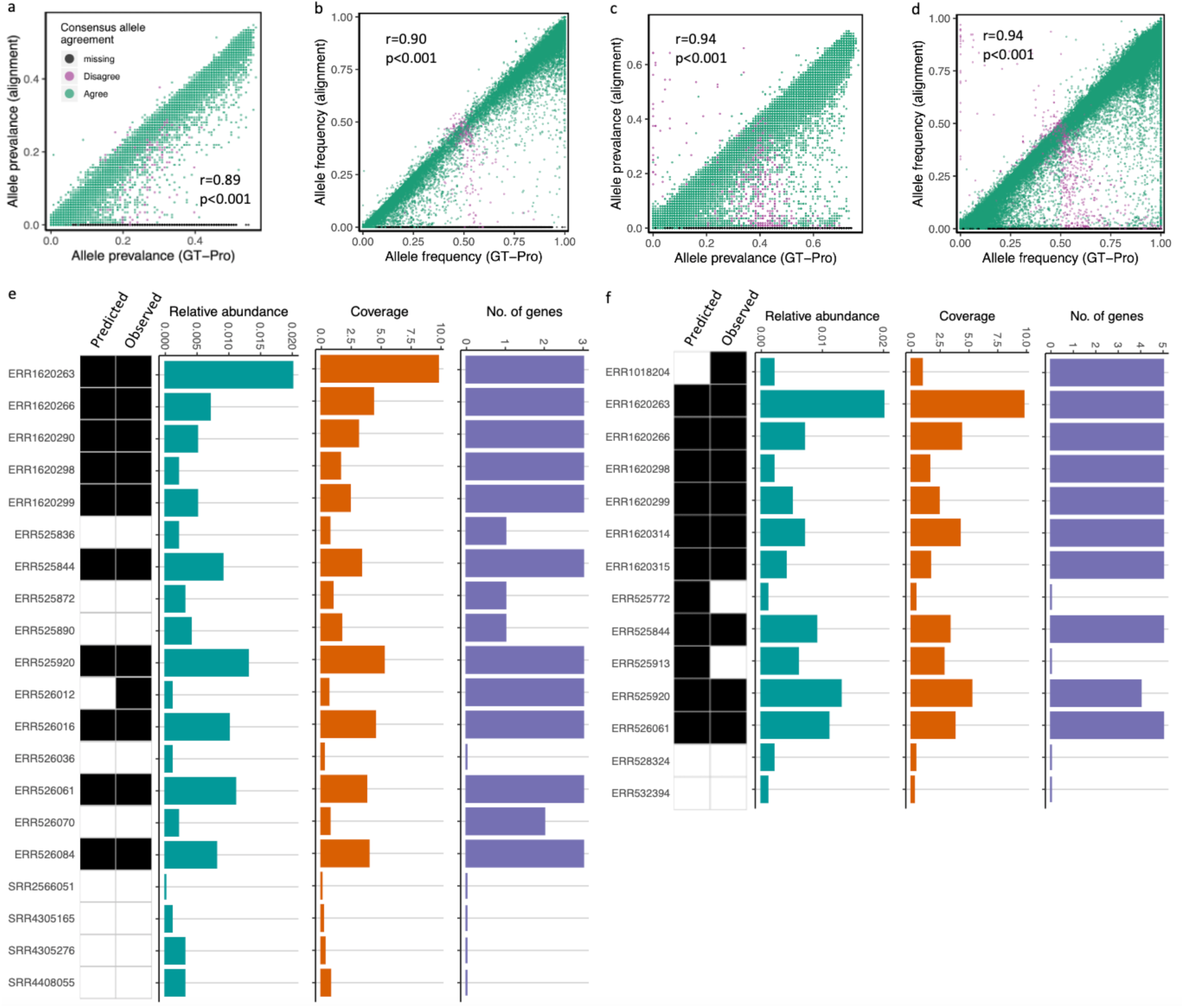
Metagenotyping and gene imputation from gut metagenomes. Comparison of metagenotypes from GT-Pro and alignment with gut microbiome samples from a North America cohort^22^ (HMP project; n=358; Table S8). As an example, we show the species *Bacteroides stercoris* (species id: 103681). Each point represents a SNP, with color indicating if the genotypes from the two methods agree (green), both methods return a genotype but the alleles disagree (purple), or only GT-Pro returns a genotype (black). Disagreements largely occur near 0.5 allele frequency, where small differences in read counts per allele can “flip” the major and minor alleles. (a) The proportion of samples in which each SNP is genotyped (prevalence) is similar with both methods. (b) Average allele frequency across samples varies across SNPs but is highly correlated between the two methods. (c and d) Comparison similar as (a and b) showing the species GCA_000431835.1 (genus: Succinivibrio, species id: 100412) from a different Madagascar cohort^16^ (n=112; Table S9). Prediction of presence/absence of *C. difficile* pathogenic gene sets in human gut metagenomes from a mix of cohorts (n=7459) (Table S14) a random forest classifier built using GT-Pro SNPs from flanking regions in 117 *C. difficile* isolates (Figures S23-S24) with 10-fold cross validation. Heatmaps show the predicted (first column) and observed (based on alignment, second column) presence (black) or absence (white) in each sample (rows). Barplots show *C. difficile* relative abundance (left), whole genome sequence coverage (middle), and number of detected genes from the pathogeneticity locus (right), all estimated by mapping reads from each sample to a *C. difficile* representative genome. Random Forest predictions correlate with abundance, coverage, and number of detected pathogenic genes (Figure S25). (l) CdtLoc genes. (m) PaLoc genes.

Next, we sought to determine if GT-Pro SNPs could be used to infer the presence of nearby genes or operons, thereby serving as biomarkers for structural variants. As a case study, we used GT-Pro SNPs in flanking genes to predict presence/absence of toxicity controlling genes in *Clostridium difficile (C. difficile)*. We used the GT-Pro SNPs from two 5’ (CD2601 and CD2602), one 3’ (trpS) gene and intergenic region to train a Random Forest classifier to predict the presence of the genomic region (CdtLoc) of three toxin genes (CD196_cdtA, CD630_cdtAB and cdtR) in a set of *C. difficile* isolate genomes downloaded independently from NCBI (Fig. S22a and S23a). In another example, we demonstrated that SNPs (cdd2, cdu1, and intergenic) flanking a pathogenicity locus (PaLoc) region could predict its presence (Fig. S22b and S23b). Next we applied these models to GT-Pro metagenomes from 7,459 samples (Fig. 5e and f). Our predictions of CdtLoc and PaLoc region presence were highly correlated with estimated presence based on read alignment to the *C. difficile* genome, especially in metagenomes where this species was more abundant, and weaker predictions were made when not all of the genes in CdtLoc or PaLoc were present (Fig. S24a and b). These results show that GT-Pro can detect structural and strain variants when they are in high LD with flanking common SNPs.

### Depicting novel and global intra-species genetic structure with GT-Pro

To evaluate the commonality of SNPs in GT-Pro database and how GT-Pro perform in metagenotying unknown metagenomes. We next used GT-Pro’s common SNPs to perform dimension reduction on the genomes in the database as well as metagenomic samples from a North American IBD cohort^24^ (n=220; Table S11) that did not contribute genomes to the GT-Pro database. Looking for evidence of subspecies genetic structure, we observed that for most species the metagenomes clustered with the genomes (Fig. 6a and b), suggesting that GT-Pro’s database represents the common diversity across diverse metagenomes. For a few species, however, we observed clusters comprised only of metagenomes (Fig. 6c and d), demonstrating that novel subspecies genetic structure can be discovered using GT-Pro common SNPs.

**Figure 6.**
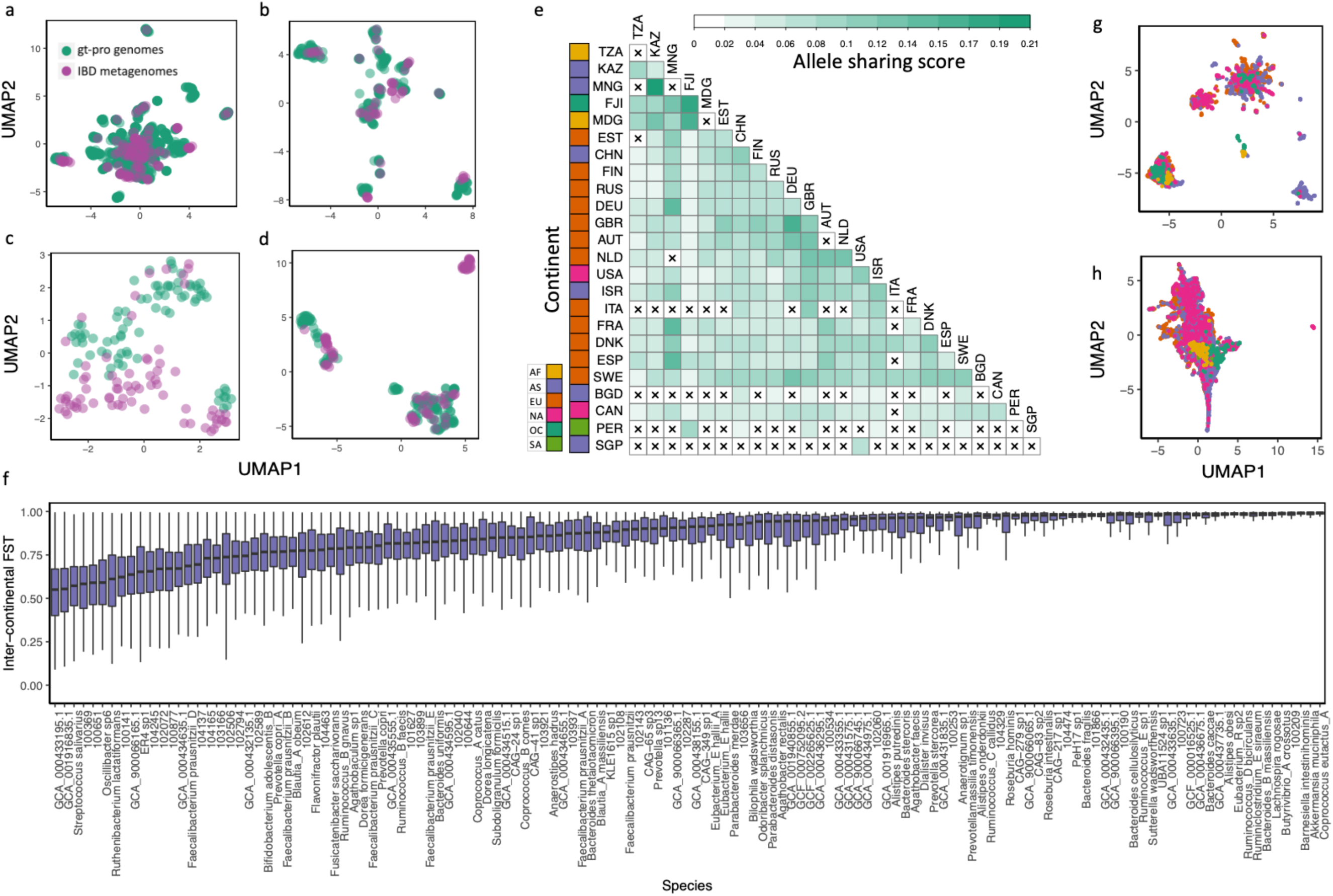
Global genetic structure in 6,452 human gut metagenomes. (a-d) Gut species differ in the amount of common SNP genetic diversity already present in sequenced genomes. Metagenomic samples from a North American IBD cohort^24^ (n=220; Table S11) (purple) are visualized in two dimensions alongside the UHGG genomes (green). Each plot is the result of applying UMAP to a matrix of genotypes at GT-Pro SNPs for one species. Each dot represents a strain of that species (major allele for heterozygous metagenomes); those closer together in UMAP space have more similar genotypes. (a) *Anaerostipes hadrus* (species id: 102528) and (b) *Ruminococcus_B faecis* (species id: 100249) are species where metagenomes lie within the diversity previously captured by genomes. (c) *Blautia_A obeum* (species id: 100212) and (d) *Dialister invisus* (species id: 104158) are species where metagenomes harbor combinations of common SNPs outside the range present in genomes, which may represent novel subspecies. (e) Heatmap of mean allele sharing scores over all species between metagenomes from different pairs of countries. Crossed cells indicates missing scores due to insufficient (< 5000) pairs of samples. (f) Analysis of inter-continental population differentiation (FST) for 78 prevalent species. Each boxplot represents a distribution of inter-continental FST for one species, ordered by medians. (g) An example of geographic patterns captured by within-species genetic variation in the GT-Pro metagenotypes of specific species. Each dot is a metagenomic sample, colored by continents. Dimension reduction and visualization performed with UMAP. The example is from *Agathobacter rectalis* (species id: 102492). Nearby samples in UMAP space have similar abundance profiles; the absence of distinct groups indicates that relative abundance does not show strong geographic clustering. (h) UMAP analysis based on the relative abundances of the 881 GT-Pro species in the same samples as (i).

Having shown GT-Pro is faster and at least as accurate as alignment-based methods for genotyping common SNPs from metagenomes, we leveraged GT-Pro metagenotypes to conduct the most geographically diverse intra-species genetic variation meta-analysis to date, encompassing 51.8 million SNPs for 881 species found in 7,459 gut samples from 31 locations across six continents (Table S13). Consistent with prior studies^4,6,9^, we observed much less allele sharing between hosts (median=0.03, IQR=0.05) than within a host over time (median=0.38, IQR=0.4), and that intra-host allele sharing varies greatly between species and hosts (Fig. S25). Inter-host allele sharing differed across countries and continents (Fig. S26a and b), generally decreasing with geographic distance (Fig. S26a and b and S27a and b) and varying across species (Fig. S28). Our results also show clear associations with degree of industrialization as well as relatedness of hosts (e.g., hosts within villages in Fiji share more alleles than unrelated hosts in North American cities) (Fig. 6e). To identify gut species with high levels of inter-continental population differentiation, we calculated FST for 78 prevalent and well-detected (see methods) species and observed large differences in the degree of differentiation across species (Fig. 6f). Species with high FST show distinct clusters of hosts, some but not all of which correlate with geography (Fig 6g), consistent with lifestyle and environment playing a role in which strains colonize a host. In contrast, hosts do not cluster as clearly based on species relative abundance (Fig 6h), emphasizing that metagenotypes may reveal microbiome-host associations missed in abundance analyses.

## Discussion

Here, we greatly extended the gut microbiome genomic variation landscape by identifying more than 100 million common core-genome SNPs from 909 bacterial species. As our solution to the bioinformatics challenge of metagenotyping, GT-Pro avoids computationally costly alignment and overcomes computing barriers. It performs strain-level analyses of microbiomes with improved accuracy, especially for low coverage species. Studies of microbiome genetic variation on a laptop or at the scale of human genome-wide association studies will be computationally feasible with GT-Pro.

It should be noted that our method comes with several limitations. First, the GT-Pro database does not capture all human gut microbial diversity. While we used 909 species, we could not use the majority of the UHGG species due to limited availability of high-quality genomes. Second, GT-Pro is analogous to a genotyping array and hence does not identify novel SNPs, which require other methods, such as alignment-based SNP calling or single-cell genome sequencing. For some species, the common SNP pool is expected to expand through additional genome sequencing. Third, a small number of species lacked species-specific sck-mers due to the presence of highly related species in the genome collection. Separate strategies such as using longer k-mers or less common SNPs could enable GT-Pro metagenotyping for these species. Fourth, although we were very selective in the choice of genomes and SNPs used for building GT-Pro, it is impossible to exclude all imperfections (e.g. incompleteness, contaminations and species misclassification) in the genome assemblies that could contribute to false SNP calls. Finally, GT-Pro does not directly genotype structural variants, which contribute significantly to intra-species genetic diversity^25^. However, we did show that GT-Pro can be used to impute insertions and deletions in high LD with common SNPs. Despite these caveats, we showed that the GT-Pro framework is general, accurate and sensitive for identifying genetic variation in metagenomes.

We envision several directions for future work. First, this study applied the GT-Pro approach to human gut prokaryotic species, and the framework could easily be expanded to other kingdoms and environments. Another extension is to develop alignment-free metagenotyping for short indels and structural variants. This study barely scratches the surface in terms of interpreting microbiome genetic variation. Towards leveraging microbiomes in precision medicine, it will be critical to comprehensively identify SNPs that are associated with disease and other traits (e.g. pathogenicity, antimicrobial resistance, drug degradation). We anticipate that GT-Pro will also be useful for detecting contamination, recombination, and horizontal gene transfer events, as well as tracking variants or strains over time, host lifestyle and geography.

## Notes

### Competing Interest Statement

The authors have declared no competing interest.

### Summary of Updates

Adding supplementary text, figures and tables.

