## Supplementary text and figure 1-28 for "Ultra-rapid metagenotyping of the human gut microbiome"

### Online methods (supplementary text) for ”Ultra-rapid metagenotyping of the human gut microbiome”

#### Methods

*Whole genome sequences and species*

To implement the GT-Pro approach for human stool metagenomics, we performed single nucleotide polymorphism (SNP) discovery using the Unified Human Gastrointestinal Genomes[1] (UHGG), a collection of 286,997 microbial genomes. All genome sequences used in the study were downloaded from the UHGG at http://ftp.ebi.ac.uk/pub/databases/metagenomics/mgnify_genomes as of September 2019. We chose the UHGG database as our major genome source for four reasons: 1) the UHGG is by-far the largest collection of gut microbial whole genome sequences, 2) microbial species in the UHGG were defined over its entire set of genomes and are well annotated both phylogenetically and functionally, 3) the genome sequences in the UHGG are originally from both isolate assemblies and metagenome assembled genomes (MAGs), and 4) the UHGG excluded genome sequences that could not be verified as coming from the human gut. The inclusion of MAGs from diverse human populations and geographic locations is critical for capturing natural genetic variation within human gut species. Since we focused on common SNPs in prevalent genomic sites (see below), sequencing and assembly errors in MAGs were not likely to get called as SNPs.

To ensure reliable SNP calling, we limited our species pool to a subset of 909 species, each with at least 10 high-quality (completeness >= 90% and contamination rate[2] <= 5%) whole genome sequences. The selected species have a median of 35 genomes. Twenty-nine species with more than 1000 genomes were further downsized to only the top 1000 genomes in terms of quality metrics and genome length. The resulting 112,904 high-quality genomes represent a large majority (>76 %) of high-quality UHGG genomes but < 20% of the total number of UHGG species. As more human gut isolate genomes and MAGs are sequenced, GT-Pro can be applied to expand species coverage.

*Species representative genome identification*

For each species, we performed whole-genome alignment among all pairs of conspecific genomes using MUMmer[3] (version 4.0.0beta2) with the default parameters. Unreliable and repeat-induced alignments were removed using the delta-filter program from MUMmer with options ‘-q -r’ and the remaining alignments were then extracted using the show-coords program with default parameters. The pairwise similarity between two genomes was calculated as the total number of matched bases using the following formula:

$$\textit{Overall similarity}= \sum_{i=1}^{n} L_i \ * \ ANI_i,$$

where $L$ is the alignment length, $ANI$ is the average nucleotide identity and $i$ represent the i’th alignment. The centroid genome with the highest average similarity to all others was selected as the representative genome for the species. This is a different approach from the UHGG study, which picked the representative genome based on the genome quality metrics only. The resulting GT-Pro representative genomes have <20% overlap (n=167) with the UHGG ones, but nonetheless have high genome quality (Figure S2).

*Genomic variation in gut microbial species*

For each species, we called SNPs by aligning each conspecific genome to the representative genome using MUMmer (same parameters and filters as above). To ensure reliable SNP calling, we removed poorly aligned regions (alignment ANI < 95% of whole-genome ANI). We then concatenated all qualified alignments and called SNPs as sites with the following characteristics: 1) two or more nucleotides, 2) present in >= 90% of genomes, and 3) minor allele frequency >= 1%. We classified all SNPs as bi-, tri- or quad-allelic and focused on bi-allelic SNPs for the subsequent method implementation and downstream analyses unless otherwise mentioned. Bi-allelic SNPs were classified as protein coding or non-coding, and coding SNPs were annotated as 1) synonymous or non-synonymous, and 2) non-sense or stop codon disruptions. We calculate the genome-wide ratio of non-synonymous to synonymous polymorphism rates (pN/pS) for each species and its expected value under the assumption that mutations occur equally across all sites, using a codon-by-codon counting method described previously[4].

*SNP linkage disequilibrium (LD) blocks*

One direct benefit of calling SNPs from high-quality genomes (versus in metagenomic reads) is the ability to accurately infer haplotypes. We scanned the reference genome of each species for clusters of SNPs in strong linkage disequilibrium (LD; $r^2 > 0.81$). We only calculated pairwise LD for bi-allelic SNPs within 10Kb sliding windows for two main reasons: 1) more efficient computation and less memory consumption and 2) LD decays below our threshold within most 10Kb windows. We calculated LD as a squared correlation for all pairs of binary-encoded SNPs for each species using PLINK[5] (version 1.9) with the following parameters: --r2, --allow-extra-chr, --ld-window 10,000, --ld-window-kb 1 and --ld-window-r2 0.81. Next, we clustered SNPs using a single-linkage strategy that is similar to the UCLUST[6] operational taxonomic unit (OTU) clustering algorithm. Briefly, we iterate through all SNPs in a window, for each SNP determining whether it should be assigned to any existing cluster based on a single-linkage strategy. Namely, if a SNP is in strong LD with one or more SNPs from a cluster, it will be assigned to that cluster. If a SNP is in strong LD with members from more than one cluster, the algorithm will treat the SNP as the link between clusters and will merge those clusters. The very first SNP or any SNP that cannot be assigned to an existing cluster starts a new cluster.

*Exaction of SNP covering k-mers (sck-mers)*

One of the main goals of this study is to leverage this SNP catalog to rapidly genotype metagenomes using short, unique genomic regions (k-mers) as makers or probes for detecting alleles that distinguish highly similar genomes from the same species in metagenomic sequencing reads. We implemented this idea by identifying any 31-base SNP-covering k-mers (sck-mers) that cover (in any of the 31 bases) each of the SNP alleles while not exactly matching other species. We chose 31 for k because 1) 31-mers can be encoded with 64-bit integers (up to 32-mer) for computational efficiency, 2) k-mers with k >= 20 are fairly unique across species[7] and 31-mers are only less unique than 32-mers in the range of choices, 3) we prefer an odd k to avoid palindrome k-mers for the ease of processing and 4) 31-mers are slightly more storage-efficient than 32-mers allowing us to store the sequences and essential annotations in 64-bit vectors. For each SNP we first extracted all possible 31-mers containing the SNP site from representative genome (sck-mers for reference allele). Next, we extracted sck-mers with the alternative allele by sliding a 31-bp window across SNP site in the multiple sequence alignment, selecting the most frequent 31-mer at each position. We then retrieved the reverse complements of all sck-mers. In this way, for every SNP site there will be up to 62 sck-mers targeting the reference or alternative allele.

*Selection of species-specific sck-mers*

To increase the specificity of genotyping with sck-mers, we removed any sck-mer shared by two or more species. For every species we generated and pooled together all possible unique 31-mers using KMC[8] (version 3.1.1) with the whole genome sequences and the options "-k31", "-m32", "-t12" and "-ci1". We then searched the sck-mers per species against the 31-mer pool from every other species. The sck-mers with hits against other species were excluded, which are a large fraction of sck-mers, especially for species with close relatives in the GT-Pro database. We required sck-mers for both alleles of a SNP.

*Identification of tag SNPs*

We identified a tag SNP for each LD block. Amongst all SNPs covered by species-specific sck-mers, we made the tag SNP the one with the most linkages ($r^2 > 0.81$) to other SNPs in the block. We estimated signs (+/-) of LD between a tag SNP and others with the Phi coefficient, adding a pseudocount of 1 to avoid divisions by zero.

*Phylogenetic resolution analysis with 16S rRNA sequences and SNPs*

We performed phylogenetic resolution analysis to assess how much within-species genetic variation was captured by different subsets of genomic sites. We compared whole-genome ANI (gold standard) to cophenetic distance based on 16S rRNA genes, 40 universal single-copy phylogenetic marker genes[9], all GT-Pro SNPs, and GT-Pro tag SNPs. We predicted 16S rRNA genes for all genomes using Barrnap (https://github.com/tseemann/barrnap) (version 0.9) with the option "-k bac". For the genomes with multiple predicted copies of 16S, we selected the highest frequency gene, or the longest ones to break ties. We predicted coding gene sequences for all genomes using Prodigal (version 2.6.3). We then searched these sequences against 40 universal single-copy phylogenetic marker genes with fetchMG (version v1.0; http://vm-lux.embl.de/~mende/fetchMG/about.html) with default options and kept output sequences as phylogenetic marker gene sequences. For each species the selected 16S and phylogenetic marker gene sequences were pooled separately for multiple sequence alignment (MSA) using Clustal Omega[10] (version 1.2.4). We concatenated the SNPs per genome into a sequence, replacing missing alleles with "." and grouping the sequences by species to obtain an MSA equivalent. We used FastTree2 to build phylogenetic trees from the MSAs for 16S, PhyEco and SNPs. Phylogenetic distances were extracted from these trees using the function cophenetic.phylo from the ape package (version 5.3) in R. We used spearman rank correlation to assess the concordance between the estimated phylogenetic distances and whole genome ANIs.

*Construction and optimization of the GT-Pro database*

We sought to efficient store a total of 2.8 billion species-specific sck-mers that cover 52.8 million bi-allelic SNPs across 881 species. For the initial optimization, we used 64-bit integers to represent the sck-mers with 00 for 'A', 01 for 'C', 10 for 'G' and 11 for 'T' and discarded the sck-mers with wildcards (e.g. 'N'). The GT-Pro database was made 4 times smaller by this step. But it was still difficult to handle, especially on a personal computer. Since for each SNP site there are up to 124 sck-mers (31 per allele type and sequence orientation), there is a considerable amount of redundant information from the following aspects: 1) two adjacent sck-mers could share 30 base overlap, 2) one pair of sck-mers could share up to 30 identical bases and only differ at the SNP site, and 3) one sck-mer itself can be used for inferring its reverse complementary version without information loss and *vice versa*. These facts guided our further optimization.

One key innovation was to not store individual sck-mers, but instead store the 61-mer centered at a given allele of each SNP and containing all of its possible sck-mers (sc-spans). For every SNP we extract two sc-spans, one for the reference allele and the other for the alternative allele. To store the sc-spans from different species, we associated additional identity information with each span, including a species identifier (6 decimal digits), major/minor allele indicator (0 or 1), and genomic coordinate of the SNP (up to 7 decimal digits) and organized these in a binary-encoded table. Although rare, approximately 1.8 percent of the sc-spans have multiple SNP sites; in one extreme case, a total of 11 SNPs were found on one sc-span. We introduced additional sc-span entries to resolve each single SNP in those cases (Figure S14). Then we separately stored in a second table the minimal information needed to recover every sck-mer from the corresponding sc-span: a 5-bit SNP offset identifying the SNP position within the sck-mer and a 27-bit index pointing to the exact entry in the sc-span table. During the lookup, the sck-mer's entire sequence can be easily recovered through the formula:

sck-mer_seq := substr(sc-span, 30 – snp_offset, 31)

and then the reverse complementary version can be inferred. To this end, the GT-Pro database after optimization consists of two tables: 1) a 10.6 GB table of sck-mers containing a 4-byte entry for each kmer and 2) a 2.4 GB table of sc-spans containing a 24-byte entry for each allele. The total storage required is 13 GB – twice as compact compared to bzip2 compression.

*GT-Pro development*

We implemented a k-mer exact matching algorithm for metagenotyping for several reasons. First, k-mers can uniquely identify SNP sites in metagenomic reads. Second, exact matching is highly efficient to compute since it can be done in a single operation at the bit level. Third, this generates highly reproducible metagenotyping results compared with other fast search methods, such as fuzzy match or probabilistic algorithms. Fourth, it is flexible to customize by adding or dropping k-mers.

For each input metagenomic read, GT-Pro first breaks it down into 31-mers and encodes them into 64-bit integers. Our goal is to find exact matches of these 31-mers with sck-mers. We take advantage of two ideas. First, a Bloom filter (*l*-filter) can be used to efficiently discard sck-mers that cannot match a query k-mer (from a metagenomic read), because they do not start with the read's length *l* prefix (first *l* letters of the k-mer, $l<k$). Second, if the sck-mer table is colex sorted, sck-mers that end with a suffix *s* of length *m* will occupy consecutive entries. Hence, we built an *m*-index to quickly locate all sck-mers that share a given suffix by simply pointing to the first and last entries for each suffix. Implementing these strategies, the following exact-match algorithm determines whether a metagenomic 31-mer query hits the database: 1) look up query prefix with *l-filter*, if found, 2) look up query suffix in the m-*mer* index, if found again, 3) examine all sck-mer entries identified by the *m-mer* index one by one and report exact matches (Figure S16). We selected $l$=32 and $m$=36 as default parameters based on the best performance in our evaluation (Figure 3). A range of values for these parameters are supported, and the optimal choices will depend on system configuration, especially the available amount of RAM.

When counting alleles for each SNP, GT-Pro enforces the following rules to reduce counting bias: 1) each read will be only counted once for one SNP, 2) if a 31-mer covering more than one SNP matches a read, both SNPs increase their count, and 3) one single count will be assigned to each SNP if a read matches multiple SNPs. For each SNP per species, GT-Pro generates an output count for both reference and alternative alleles. Species abundance is estimated by first summing reference and alternative allele count into a SNP count and then taking the average SNP count across entire genome per species.

GT-Pro uses a concise table-shaped format for its output, in which every row represents a bi-allelic SNP site. Each row has exactly 8 fields: species, SNP ID, contig, contig position, allele 1, allele 2 and coverage of allele 1 and coverage of allele 2.

*Comparing GT-Pro to alignment method*

We compared GT-Pro to alignment-based metagenotyping in a series of simulations and data analyses designed to evaluate different aspects of computational performance and accuracy. We use two specific alignment-based methods to ensure our conclusions generalize across alignment-based strategies. Both methods implement a similar three-step genotyping paradigm, including aligning short reads to reference genomes (read mapping), piling up mapped reads for counting coverage per site (pile-up), and calling SNPs from site coverage profiles (SNP calling). One method is the metaSNV[11] software package (http://metasnv.embl.de/; September 2019), which uses BWA[12] (version 0.7.17-r1188) as the aligner, SAMtools[13] mpileup (version 1.9) for pile-up, and is distributed with a specific database (freeze9). The second method is a pipeline similar to MIDAS[14] (https://github.com/snayfach/MIDAS; September 2019) but with improved parallel processing for higher throughput. It uses Bowtie2[15] (version v2.3.2) for read mapping, PySAM (https://github.com/pysam-developers/pysam; version 0.12.0.1) for pile-up, and a database generated with tools from the MIDAS package using the representative genomes from GT-Pro).

Unless otherwise noted, we ran GT-Pro with the full database and default parameters and removed from output the sites with only single site coverage. We ran metaSNV with default database (freeze9) and parameters. We ran our bowtie pipeline with the aforementioned database and default parameters except a minimum site coverage of 5 instead of 10, which is adjusted to be consistent with metaSNV. For all paired-end samples, we processed only fastq 1 samples for the simplicity of comparison and analysis.

*Computing performance evaluation environment*

We evaluated performance on two typical computing environments: server and laptop. For server-based evaluation we set up an AWS EC2 instance with the following specifications: AWS r5.12xlarge, 24 physical CPU cores (48 vCPU), Intel 8175M CPU @ 2.50GHz, 384 GB RAM and EBS gp2 RAID array providing 780 MB/sec bandwidth. The laptop environment was on an Apple MacBook Pro (15-inch, 2019) with the following specifications: 8 physical CPU cores (16 vCPU), Intel(R) Core(TM) i9-9980HK CPU @ 2.40GHz, 32 GB 2400 MHz DDR4 RAM and 2TB APPLE SSD AP2048M. For each environment, we measured both speed and peak RAM use for each method. We did so for GT-Pro while tuning the values of l and m - two parameters that are derived from the l-bit and m-bit filter.

*Accurate identification of SNPs from simulated metagenomes*

To evaluate the accuracy of metagenotyping by GT-Pro and to compare it to alignment-based metagenotyping, we conducted a series of simulations that used reads generated from whole genome sequences. We used InSilicoSeq[16] (version 1.4.2) with the options "--model HiSeq" and "--n_reads 2000000" to simulate reads with Illumina length and error characteristics. This generates two paired-end read files each containing ~1 million 126bp-long reads from each genome. For simplicity, we proceeded with only the forward reads. We varied genome coverage across simulations by randomly drawing reads from the simulated metagenomes. The number of reads required for a level of coverage c was estimated by the following formula: number of reads = c * genome length / 126. For example, to provide a 10x coverage for a genome with a size of 5M bp, a rough number of 396,825 (10 * 5,000,000 / 126) 126bp-long reads is needed. We used eight levels of coverage: 0.001x, 0.01x, 0.1x, 1x, 2x, 5x, 10x and 15x.

We used isolate genomes (n=978) from the Culturable Genome Reference (CGR) study for these simulations. These genomes were also included in the UHGG genome collection under the label of "BGI" and covered in GT-Pro database. These genomes were cultivated from fecal samples of healthy humans and characterized as non-redundant and high-quality draft genomes. To ensure GT-Pro does not gain an advantage due to the fact that the genomes used for simulation were also included in the UHGG genome collection and contributed SNPs to GT-Pro database, we removed information from these isolates from the GT-Pro database by excluding the isolates and rebuilding the database from scratch for the purpose of these simulations.

First, we simulated a simple scenario where each metagenome only contained reads from a single isolate. We selected a total of 232 isolate genomes from 152 species with at least 5000 SNPs covered in the GT-Pro database. Each isolate genome was randomly assigned a sequencing coverage from the eight levels. To evaluate metagenotyping accuracy, we identified SNPs using whole-genome alignment between each genome used to simulated reads and the GT-Pro reference genome for that species. We additionally imputed genotypes within LD blocks by assigning the reference allele if the tag SNP has the reference allele, and the alternative allele if the tag SNP has the alternative allele. SNPs outside LD blocks and SNPs with Phi<0 with their tag SNP were not imputed.

In a second scenario, we generated sequencing datasets containing reads from a pair of conspecific genomes to simulate metagenomic samples with strain mixtures. We used a total of 80 genome pairs, each from a distinct species. For each pair, we generated 8 metagenomic samples by fixing the sequencing coverage for one genome at 15x and generating reads for the other genome with each of the eight coverage levels. In this way, we generated a total of 640 metagenomic samples from 80 species with the coverage ratio of a strain 1 versus strain 2 ranging from 0.001:15 to 15:15. To evaluate accuracy, we called SNPs in the whole-genome alignment of the two strains used to simulate reads to each other and to the reference genome. Based on the alignments, each GT-Pro SNP was labeled as homozygous reference allele, heterozygous, or homozygous alternative allele.

We ran GT-Pro and our bowtie-based alignment pipeline on the simulated reads from each simulation. No minimum site coverage or any other filtering criteria were applied to either methods here in order to minimize the biases that could be introduced by this preprocessing.

Next, we compared metagenotypes to the known values and estimated various performance measures. Our evaluations focused on correct identification of heterozygous SNPs across coverage ratios and on the accuracy of the alleles in the metagenotypes, as well as accurate estimation of species and strain abundance using SNP site coverage and our ability to use tag SNPs to impute genotypes of other GT-Pro SNPs. Since we separately estimated the genetic diversity that can be captured by GT-Pro SNPs in genomes (see above), these evaluations focused on correct allele calls at GT-Pro SNP sites (i.e., we only included SNPs that are potentially able to be genotyped by both GT-Pro and our bowtie-based pipeline).

In simulations with a single species per metagenome, false positives (FP) are incorrect genotypes as well as any genotypes called for species other than the one from which reads were generated. True positives (TP) are correct genotypes in the species from which the reads were generated. Hence, we can efficiently compute the false discovery rate (FDR) as 1 minus the ratio between the sum of TP sites and the sum of all reported sites. False negatives (FN) are sites with no genotype or an incorrect genotype, so sensitivity can be calculated as TP/(TP+FN) or the ratio of TP sites to all GT-Pro sites.

In simulations with strain mixtures, we focus on genotypes reported for the species from which reads were simulated (ignoring other species). We estimate FDR and sensitivity for calling heterozygosity (i.e., two genomes have different alleles), as well as accuracy of genotypes (which can be homozygous reference, heterozygous or homozygous alternative). At homozygous sites, FP are heterozygous calls and homozygous calls with the incorrect allele.

*Performance evaluation on publicly available metagenomes*

The sequencing samples used for performance evaluation were human gut microbiome fecal samples from Hadza hunter-gatherers in Tanzania[17], which are available at https://www.ncbi.nlm.nih.gov/ with accession ID SRP110665. The selection of samples and reads is arbitrary. For each baseline method, the chosen 36 samples were processed in their entirety. For GT-Pro laptop, each sample contributed just its first 1.5 million reads. For GT-Pro server, the first and the last 1.5 million reads were used. We used sar and htop to monitor CPU and I/O utilization during server runs.

*Metagenotyping 25,133 human microbiomes with GT-Pro*

To further demonstrate the computational efficiency of gt-pro, we applied GT-Pro to metagenotype a total of 25,133 publicly available human metagenomes from diverse body sites, biogeographic regions, lifestyles and age groups (Table S5 and S6). These samples comprise >40 trillion bases in >400 billion reads from 45,683 fastq files. We used two AWS EC2 instances (type: c5.18xlarge) for processing the samples, half on each instance. Each instance has 72 vCPUs, 144 GB RAM, 25 Gbps network bandwidth and 19 Gbps EBS bandwidth. Due to the limits of EBS storage space, we wrote a Bash script for processing the samples in small batches with four steps: 1. download SRA files with accession numbers, 2. extract fastq files from SRA files, 3. run GT-Pro on fastq files and 4. deposit metagenotyping results to S3 storage and clean out EBS space. It took less than three weeks to process these samples. We estimate that the same task would take more than a year on the same system using alignment-based metagenotyping. We observed that gt-pro shifts the computational bottleneck from genotyping reads to data downloading, transferring and decompression, which were responsible for two thirds of its processing time.

*Accurate metagenotyping from gut metagenomes*

To further evaluate GT-Pro versus alignment-based genotyping on more complex communities, we metagenotyped gut metagenomes from 4 cohorts (Table S7-10) from Tanzania[17] (N=40), North America[18] (HMP; N=358), Madagascar[19] (N=112) and Sweden[20] (mother-infant pairs; N=400). We choose diverse samples to explore the applicability of GT-Pro to human gut microbiomes across populations. With the Tanzania cohort, we compared GT-Pro to metaSNV to evaluate how the two different databases affect metagenotypes. From the output of both methods, we count the number of species, total number of sites genotyped of all species, and median number of sites genotyped per species. With the HMP and Madagascar cohorts, we compared GT-Pro to our bowtie-based pipeline using GT-Pro reference genomes (i.e., same genome database) to isolate the effect of k-mer exact matching versus alignment. We compared within-sample heterozygosity and across-sample allele presence and frequency between the two methods, using only GT-Pro SNP sites (i.e., sites that can theoretically be genotyped by both methods, as in simulations). The methods were compared using Jaccard index on genotyped sites and Pearson correlation of genotypes. Jaccard distance was calculated as the number of genotypes (major allele) that agreed by both methods divided by the number of SNP sites genotyped by both methods. Pearson correlation coefficient is calculated using the allele frequencies at sites genotyped by both methods. The consensus allele is identified per SNP site as the genotype with the higher across-sample allele presence. These statistics were computed using genotypes from all species per sample and separately for each species. With Swedish mother-infant cohort, we compared pairwise Jaccard distances between samples using all reported genotypes per species (i.e., here the alignment-based method could use SNPs that are not covered by the GT-Pro database).

*Flanking genes analysis of Clostridium difficile pathogenicity locus*

Our goal of this analysis is to test if GT-Pro can predict structural variants using flanking GT-Pro SNPs. We used the medically important presence/absence of genomic regions containing toxin genes in *C. difficile* strains. In one example, we used SNPs from two 5' (CD2601 and CD2602) and one 3' flanking gene (trpS) to predict the presence/absence of the CDT locus (CdtLoc) which contains the binary toxin encoding gene cdtAB and a response regulator gene cdtR (Figure S22a). Second, we used SNPs from two genes (cdd1 and cdu1) as well as the regions in between that flank the pathogenicity locus (PaLoc; contains five toxin genes tcdA, tcdB, tcdC, tcdE, tcdR) to predict the presence/absence of this region (Figure S22b).. In contrast to the PaLoc region which has several different known structural variants, the CdtLoc region only has one known deletion where a unique 68-bp sequence replaces cdtAB and trpS.

We downloaded 1171 *C. difficile* genomes deposited in NCBI RefSeq (ftp://ftp.ncbi.nlm.nih.gov/genomes/refseq/bacteria/Clostridioides_difficile/) as of May 2018. We used checkM[2] (version 1.0.11) to estimate the genome quality. We removed the genomes with completeness < 0.97, contamination > 0.02, strain heterogeneity > 0.25, GC content > 30% or genome size > 4.6 Mbps. We also used dRep[21] (version 1.4.3) to remove highly similar genomes, and any genomes with ANI cutoff > 0.99 and minimum coverage of the larger genome > 0.95 to a non-redundant genome were removed. This resulted in 114 high-quality, non-redundant *C. difficile* genomes. Amongst these, we proceeded to analyze 110 where the flanking genes for both toxin loci were present.

We annotated the CdtLoc and PaLoc regions in the *C. difficile* genomes using BLAST[22] (version 2.7.1), requiring both >20% horizontal coverage and >90% sequence identity to call hits to each toxin gene. We found that this subset of 110 *C. difficile* strains contains a variety of deletions in the PaLoc region, which could result from natural structural variants or incomplete genome assemblies. In contrast, 15 strains harbored the known 68-bp substitution in CdtLoc and the rest had the complete locus.

Next, we investigated what GT-Pro SNPs are found in and flanking CdtLoc and PaLoc. We found no SNPs in the CdtLoc region (confirmed by alignment of all sck-mers to the *C. difficile* genomes), but we identified 180 SNPs that locate in the flanking genes: 62 in CD2601, 60 in CD2602, 34 in trpS and 24 in inter-genic regions (Figure S22a). For the PaLoc region, we identified 56 SNPs in the flanking regions: 7 in cdd1, 32 in cdu1, and 17 in between the two genes (Figure S22b). We removed SNPs with the same genotype in all genomes because they have no power to differentiate genomes for the purpose of classification and kept 148 and 40 SNPs for the CdtLoc and PaLoc region, respectively.

To test whether these GT-Pro SNPs in flanking regions can be used to infer the structural variants, we built a random forest classifier for the presence/absence of CdtLoc (and separately for PaLoc) using flanking SNPs as features. We label a genome toxigenic only if it has a complete PaLoc region (e.g. presence of five genes) and the partial presence of a gene as presence. To ensure classifier to correctly deal with genomes or samples with all flanking SNPs missing or absent, we added to training a mock genome with all feature SNPs and prediction label marked as absent. The random forest classifier was then trained with randomForest package (version 4.6-14) in R with default hyperparameters and run through 10-fold cross validation.  Predictive power of each SNP was determined using importance function in the same package, which quantifies the contribution of each variable to prediction accuracy by permuting it.

We screened genotype profiles produced by GT-Pro in a total of 7,459 gut samples. We found 135 and 52 samples containing one or more flanking SNPs for the CdtLoc and PaLoc region, respectively. For the simplicity of analysis, we only continued on a subset of samples for both the CdtLoc (n=41) and PaLoc (n=33) region, which contains all top 20 SNPs with highest feature importance. We extracted and binarized the flanking SNPs as feature inputs and classified samples with the aforementioned random forest classifier trained on genomes. The predicted presence/absence of the PaLoc or CdtLoc region in a sample was determined by probability present > 0.6.

For the same samples, we used an alignment method (MIDAS) to determine observed presence/absence of the CdtLoc and PaLoc region, *C. difficile* relative abundance, whole genome sequence coverage, and number of detected genes from the pathogeneticity locus. Reads from each sample were mapped to a *C. difficile* representative genome (UHGG ID: GUT_GENOME142303). The presence of a gene was determined by horizontal coverage (> 50% of full sequence length) by reads, and the observed presence of a region was determined by percentage of genes present (>= 80%), i.e. presence of the PaLoc region requires at least 4 genes present and CdtLoc requires all 3 genes present.

*Novel subspecies detection in IBD microbiomes*

To explore genetic structure within species, selected a set of metagenomic samples (n=220) from a North American IBD cohort[23]. This data is from a 2019 manuscript and did not contribute any MAGs to the UHGG collection, so their strains are likely to be at least somewhat novel compared to the genomes we used to build the GT-Pro sck-mer database. We sought to determine whether the common SNPs covered by the database could be used to identify novel allele combinations. To do so, we applied GT-Pro to the 220 metagenomes and then performed dimension reduction on the resulting metagenotypes along with the genotypes of the genomes used to build the sck-mer database. If metagenome-specific samples cluster outside the variation of genomes in the dimension reduction, it would suggest that they carry novel SNP allele combinations. A cluster of metagenomes would indicate a group of related strains that differ from known strains and could represent a new subspecies. We used the following steps for performing this exploratory analysis: 1) identified and estimated the frequency for the major allele per SNP per sample, 2) identified the major allele per SNP site as the allele presents in the most samples, 3) recorded the matrix of major allele frequency for each sample, 5) extracted a similar matrix of binary major allele presence/absence for each genome used to build the GT-Pro sck-mer database, and 6) applied UMAP analysis and plotted metagenomes and genomes in UMAP coordinates.

*Global population genetic diversity of human gut microbiomes*

From all metagenotyped samples, we selected a total of 7,459 gut samples from 31 locations across all continents except Antarctica (Table S13) for a global population genetic diversity analysis. The selection is limited to human gut samples and a single sample per host to minimize potential biases that repeated sampling might introduce.

We calculated an allele sharing score between each pair of samples as an estimate of their genetic similarity. We identified shared SNP sites as those genotyped in both samples, and then defined the allele sharing score as the number of shared SNP sites with the same major allele in both samples divided by the number of all genotyped SNP sites in the lesser sample, i.e. min(# of genotyped sites in sample 1, # of genotyped sites in sample 2). We adopted this measure of similarity mostly because many samples are very different in the number of reads.

We down-sampled 84000 pairs of samples and calculated the fixation index statistic (FST), an estimate of population differentiation, for 78 prevalent and well-detected species (present in >500 inter-continental sample pairs, each pair has >1000 shared SNP sites). To reduce bias due to disparate sample size, we performed down-sampling by randomly drawing 6000 pairs of samples per continent pair and ignored continent pairs with <6000 pairs of samples. FST is defined as 1 minus the ratio of within-sample and between-sample nucleotide diversity ($\pi$) , or more formally, $1 - (\pi 1 / 2 + \pi 2 / 2) / \pi b$, where $\pi i$ (i = 1 or 2) represents within-sample $\pi$ for sample i and $\pi b$ represents the between-sample $\pi$ between sample 1 and 2. To estimate within-sample $\pi$, we first calculate the single site $\pi$ of a genotyped SNP site as $2 * a1 * a2 / (a1 + a2) / (a1 + a2 - 1)$, where ai (i = 1 or 2) represents the coverage of the i'th genotype of a SNP site, and then take an average of single site $\pi$ across all genotyped sites. Similarly, to estimate between-sample $\pi$, we first calculate the single site $\pi$ of a shared site as $(a11 * a12 + a21 * a22) / (a11 + a21) / (a12 + a22)$, where aij (i, j = 1 or 2) denotes the coverage count of the i'th genotype of a SNP site in sample j, and then take an average of single site $\pi$ across all shared sites. This approach only accounts for bi-allelic SNPs.

*Data availability*

Genome assemblies used in this study were downloaded from the UHGG database hosted at MGNify (http://ftp.ebi.ac.uk/pub/databases/metagenomics/mgnify_genomes). The GT-Pro SNP databases we generated in this study is available in an AWS S3 bucket with public access permission (s3://jason.shi-bucket/public/gt-pro_main). The implementation and documentation of GT-Pro is available on the github (https://github.com/zjshi/gt-pro). GT-Pro itself is written in C++ and released as open-source software under the MIT license.

**Supplementary figure legends**

Figure S1. Implementation of the gtpro *in sillico* metagenotyping framework for the human gut microbiome (layout mirroring Figure 1). We identified 909 gut species with at least 10 high-quality genomes in the UHGG[1]. More than 104 million common SNPs were called through whole genome alignment of conspecific genomes. Up to 124 (31 X 4) candidate k-mers were extracted per bi-allelic SNP site, a total of 1.2 trillion candidate SNP-covering k-mers (sck-mers) were extracted, and among them 5.7 billion sck-mers were determined to be species-specific. The resulting GT-Pro database directly covers 52.8 million SNPs from 881 species. In parallel, a total of 6.8 million LD blocks were detected by clustering SNPs based on pairwise linkage disequilibrium (LD). Within LD blocks, about 15 million SNPs were assigned as tag SNPs, including standalone SNPs (not in LD blocks with any other SNP). Tag SNPs must be covered by at least one species-specific sck-mer for each allele. An additional 15.7 million "indirectly covered" SNPs without sck-mers are in strong LD (r2 > 0.81) with a tag SNP. Database characteristics are compared to those of Schlossnig et al[24]. and the 1000 genome project catalog of human genetic diversity[25].

Figure S2. Distribution of genomes used to build the GT-Pro human gut database according to their quality metrics, including (a) completeness and (b) contamination rate. The black dashed lines represent the mean from the representative genomes.

Figure S3. (a) Counts of bi-, tri- and quad-allelic common SNPs discovered in 909 genomes, summarized across gut bacterial phyla. (b) Number of SNPs per bacterial phylum colored by being in GT-Pro (green), in an LD block with a SNP in GT-Pro (orange), or not detectable by GT-Pro (purple).

Figure S4. Characteristics of common SNPs (from top to bottom) across species ordered by phylum and number of SNPs: number of discovered alleles, location of bi-allelic SNPs in coding or non-coding sequences, proportion of coding mutatinos that are synonymous versus non-synonymous, and types of non-synonymous mutation (non-sense, stop codon disruptive or non-stop codon).

Figure S5. (a and b) Correlation between the number of genomes and (a) the number of discovered SNPs or (b) the number of SNPs per Kb. The black dashed line in both (a) and (b) represents the linear regression line. (c) Comparison of average LD block size (e) across gut bacterial phyla.

Figure S6. (a) Rarefaction curves of common SNP discovery for the species with at least 50 genomes. Each curve represents a species and is made by calculating the number of SNPs that can be discovered using down-sampled subsets of genomes per species. The subsets start with 10 genomes and increase by 10 genomes up to the true number. This down-sampling was repeated 10 times and the mean number of SNPs discovered at each subset size is plotted. (b) Histogram showing the percent increase in common SNPs discovered when adding the last 10 genomes for each species in (a). Most species have a low rate of increase, indicating that SNP discovery has leveled off. But a few species have 8-10% increases, suggesting that more common SNPs remain to be discovered.

Figure S7. Correlation between the number of SNPs per Kb and the nucleotide diversity. Each dot in the plot represent a species and dot color reflects the phylum of the species. The black dashed line is the linear regression line.

Figure S8. Distribution of pN/pS per species across gut bacterial phyla. The dots above the black horizontal line represent the species with both high pN/pS (> 0.5) and low pS (< 0.007).

Figure S9. Examples of LD distance decay as genomic distance between SNPs increases: (a) Streptococcus salivarius (genus: Streptococcus) (species id: 100113), (b) Bacteroides thetaiotaomicron (genus: Bacteroides) (species 100196), (c) an unclassified species from Escherichia genus (species id: 102506) , (d) Roseburia inulinivorans (genus: Roseburia) (species id: 100271), (e) Veillonella parvula (genus: Veillonella) (species id: 101349), (f) Vibrio parahaemolyticus (genus: Vibrio) (species id: 102331), (g) Acinetobacter baumannii (genus: Acinetobacter) (species id: 102344), (h) GCA_001916965.1 (genus: COE1) (species id: 104431).

Figure S10. Number of LD blocks identified with increasing r2 cutoff. Each observation represents a species.

Figure S11. (a) Distribution of LD blocks per Kb across species, including all singleton SNPs as LD blocks. Red vertical line: one LD block per Kb. (b) Rarefaction curves of number of LD blocks detected with down-sampling genomes for all species with more than 1000 genomes.

Figure S12. Summary of average SNP covering depths for all species. SNP covering depth is the number of species-specific sck-mers for a SNP. Horizontal dashed lines and the bar on the right-side group species by average SNP covering depth, with darker red indicating more sck-mers per SNP.

Figure S13. Resolution of within-species genetic diversity captured by different subsets of SNPs. The genetic distances (phylogenetic tree branch length by FastTree 2.0) between all pairs of conspecific genomes were computed using five sets of SNPs: entire catalogue of common SNPs regardless of having sck-mers (all), all SNPs in the GT-Pro database (full), GT-Pro tag SNPs (tag), only SNPs in 40 universal marker genes[9] (marker), and only SNPs in the 16S rRNA gene (16S). (a) Distances using each subset of common SNPs (rows) compared to pairwise ANI (includes rare SNPs) for eight species (columns) with the most genomes. Species (from left to right): Agathobacter rectalis (genus Agathobacter) (species id: 102492), Bifidobacterium infantis (genus: Bifidobacterium) (species id: 101292), an unassigned species from Ruminococcus_E genus (species id: 100258), Klebsiella pneumoniae (genus: Klebsiella) (species id: 102538), Vibrio cholerae (genus: Vibrio) (species id: 102311), Salmonella enterica (genus Salmonella) (species id: 102366), Campylobacter jejuni (genus: Campylobacter) (species id: 102422), an unassigned species from Escherichia genus (species id: 102506). (b) Heatmap of Spearman's rank correlations between pairwise ANIs and the five common SNP-based distances (rows) for all 909 gut bacterial species (columns, sorted from low to high correlation). (c) Boxplot summarizing the distributions of correlations in each row of (b).

Figure S14. Examples for overlap encoding in the sc-span table. (a) An example of a sc-span entry which only has a single SNP (red). Two underscored sequences are the full-length (61bp) sc-spans for both major and minor allele. (b) An example of a sc-span entry which has two SNPs (red, blue). A third full-length sc-span (underscored and labeled as sc-span') was introduced to resolve the additional SNP without duplicating data uneccessarily by creating a span for each SNP.

Figure S15. Summary of number of SNPs (a) and sck-mers (b) in full and tag version of GT-Pro database. Each observation is one species.

Figure S16. Development and optimization of GT-Pro species-specific sck-mer data structures and algorithms.(a) Workflow for metagenotyping. In the GT-Pro database's sck-mer table, all species-specific sck-mers have been binary encoded, pooled, and sorted in Colex order. The bits of sck-mers were split into l-mer prefixes for a Bloom filter (blue) and m-mer suffixes to build an index (green) whose entries point to consecutive records in the sck-mer table (violet). Efficient exact-matching of k-mers from metagenomes starts with extracting all k-mers present in each sequencing read (input). These are passed through the l-mer Bloom filter. Only k-mers passing the filter can be exact matches to sck-mers. To determine if they are, the range of entries in the m-mer index corresponding to each l-mer hit is queried with an exact-match algorithm. Dashed black arrows indicate processes occurring during database development, and solid gray arrows indicate the metagenotyping workflow. (b) Schematic showing details of the sck-mer table data structure. (c) Details of the SNP-centered spans (sc-span) table.

Figure S17. Sensitivity of GT-Pro and alignment at homozygous sites in simulations with reads from two isolate genomes per species, across a range of simulated sequencing coverage combinations. One genome is always simulated at 15x coverage, and the less abundant genome has variable coverage. Sensitivity is the probability of correctly calling a site homozygous when the isolate genomes have the same allele at a GT-Pro SNP site.

Figure S18. Distribution across species of odds ratios quantifying agreement between tag SNPs being genotyped and non-tag SNPs in the same LD block being genotyped (using the full version of the GT-Pro database). High odds ratios indicate species where non-tag SNPs are rarely absent if the tag SNP for their LD block is present. Odds ratios are based on genotyped sites in simulations in Figure 5a.

Figure S19. (a-c) Amount of diversity captured in metagenotyping results on a Tanzanian cohort[17] (n=40; Table S7) for GT-Pro (full SNP set) versus alignment (metaSNV). Method used their own databases, which contain different species, so results reflect the combination of what theoretically can be genotyped (species for each, SNPs per species for GT-Pro) and the sensitivity of each method to call genotypes across species with different coverages in the data. (g) Total number of genotyped sites per sample. (h) Median number of genotyped sites across detected species in each sample, and (i) number of detected species per sample.

Figure S20. Agreement of metagenotyping output between GT-Pro and alignment in samples from the Human Microbiome Project[18] (HMP, above; Table S8) and Madagascar cohort[19] (MDG, below; Table S9). Agreement was measured with both Jaccard distances (left) and correlation of allele frequencies (right) at all genotyped sites per sample.

Figure S21. Heatmaps showing pairwise genetic distance (Jaccard metric) between samples from a mother-infant cohort[20] (n=400; Table S10) based on different subsets of SNPs (left to right) for a given species: all SNPs genotyped by alignment baseline method (with MIDAS, includes rare SNPs), all common SNPs genotyped by GT-Pro, and all SNPs identified by mapping reads to the 16S gene. (a) Bacteroides_B dorei (species id: 102478), detected in 75 samples with median coverage 39.4X. Subjects (n=9) with all M, 4M and 12M samples detected were used for the plot. (b) Bifidobacterium adolescentis_B (species id: 102395), detected in 51 samples with median coverage 11.9X. Subjects (n=5) with both M and 4M samples detected were used for the plot.

Figure S22. Pathogenic genes and GT-Pro SNPs in their flanking regions on contigs of the representative genome of *C. difficile* from GT-Pro database. SNPs were used to predict presence/absence of the pathogenic gene sets. Top: gene locations (colors: genic regions), bottom: flanking SNP locations (colors: genic or intergenic labels). (a) CdtLoc. (b) PaLoc, which spans two contigs.

Figure S23. Prediction of presence/absence of pathogenic gene sets in the genomes 110 *C. difficile* isolates (Table S14) with a random forest classifier built using GT-Pro SNPs from flanking regions and 10-fold cross validation. Rows: SNPs (sidebar colors indicate genic regions), columns: genomes (sidebar colors indicate pathogenic gene set presence/absence in the genomes (observed) and according to the random forest (predicted). Heatmap cells show SNP allele: reference (green), alternative (purple), not detected (light grey). The random forest yielded a perfect prediction from the cross validation. SNPs are ordered by genomic coordinate. (a) CdtLoc and (b) PaLoc.

Figure S24. Random forest model is more confident of the presence of the entire pathogenic gene set when more of the genes are detectable in the metagenome. Plotted values for each metagenome are the random forest score (predicted probability of the entire pathogenicity gene set being present based on flanking SNPs) and the number of genes with >50% of length covered at least 1 read in the metagenome. Dotted line, r2, and p-value based on linear regression model. (a) CdtLoc and (b) PaLoc.

Figure S25. Allele sharing increases with relatedness of metagenomes. Scores calculated between pairs from a set of randomly selected samples (Table S12) from different hosts (inter-individual) and different samples from the same host (intra-individual) host, and same technical replicates from the same sample.

Figure S26. (a and b) Average allele sharing increases with geographic proximity. Scores calculated between pairs of samples across cohorts from different countries within the same continent (a) and from different continents (b). Continents: Africa (AF), Asia (AS), Europe (EU), North America (NA), Oceania (OC), South America (SA).

Figure S27. Comparison of allele sharing scores between pairs of samples from (a) same versus different continent and (b) same versus different country.

Figure S28. Pairwise allele sharing scores by individual species between samples from (a) the same continent and (b) different continents. Each point represents a mean allele sharing score for a species. Black horizontal line represents pairwise allele sharing scores using all species. The species are colored by their phyla.

#### Supplementary figures

##### Figure S1

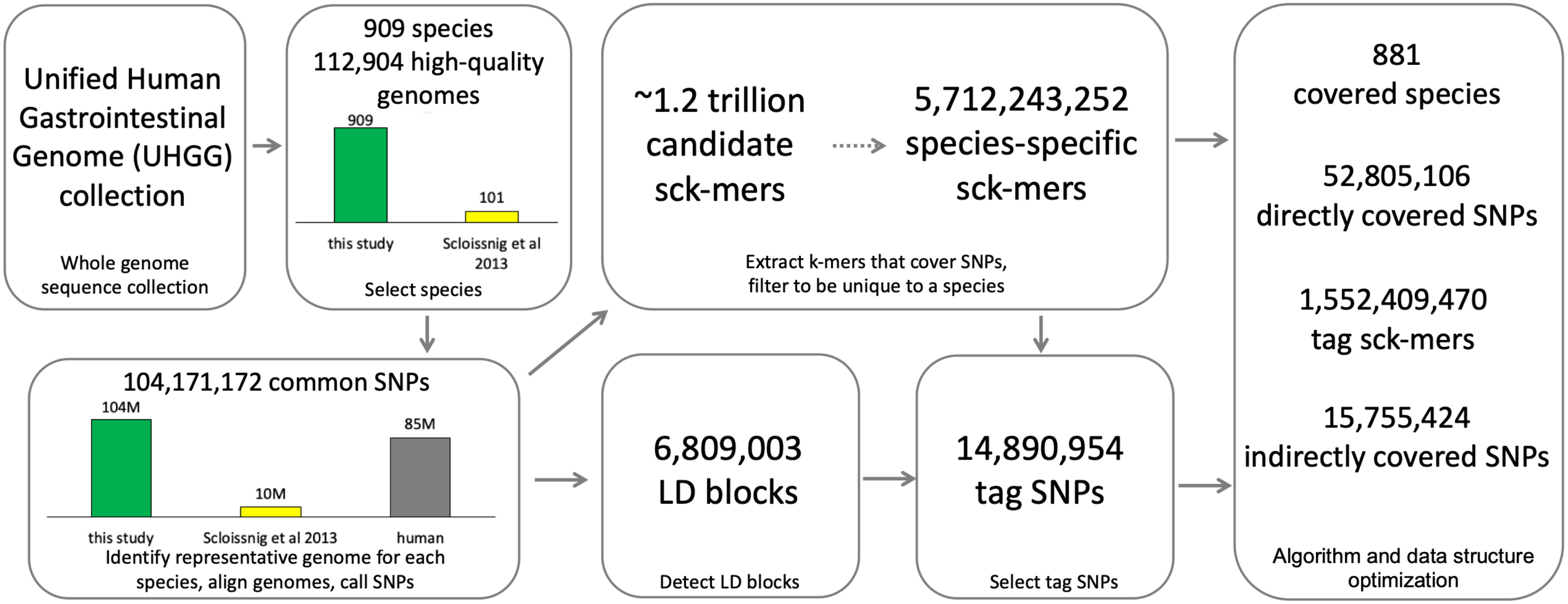

**Figure S2**

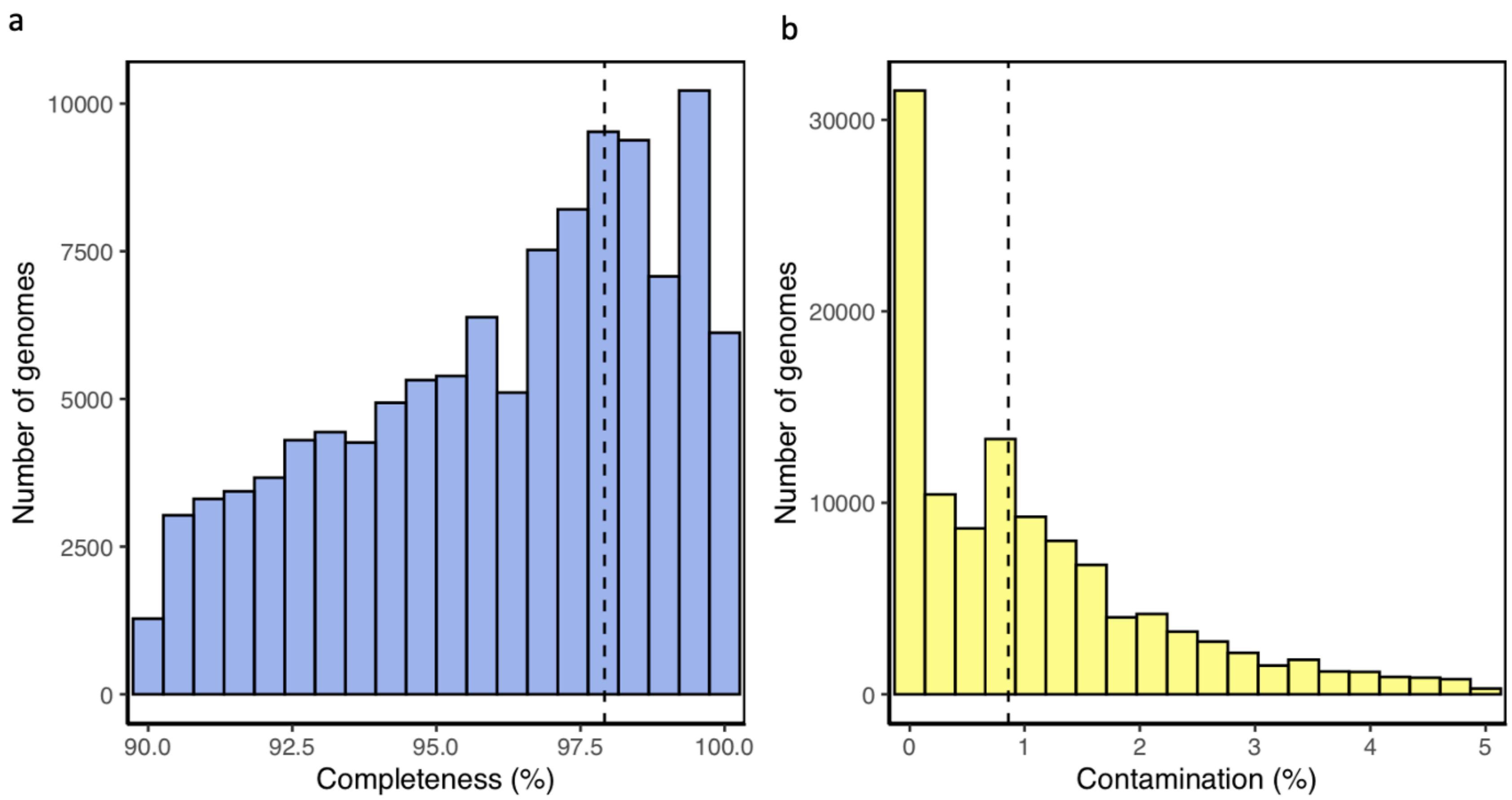
a
b
Number of genomes
10000
7500
5000
2500
0
90.0
92.5
95.0
97.5
100.0
Completeness (%)
Number of genomes
30000
20000
10000
0
0
1
2
3
4
5
Contamination (%)

**Figure S3**

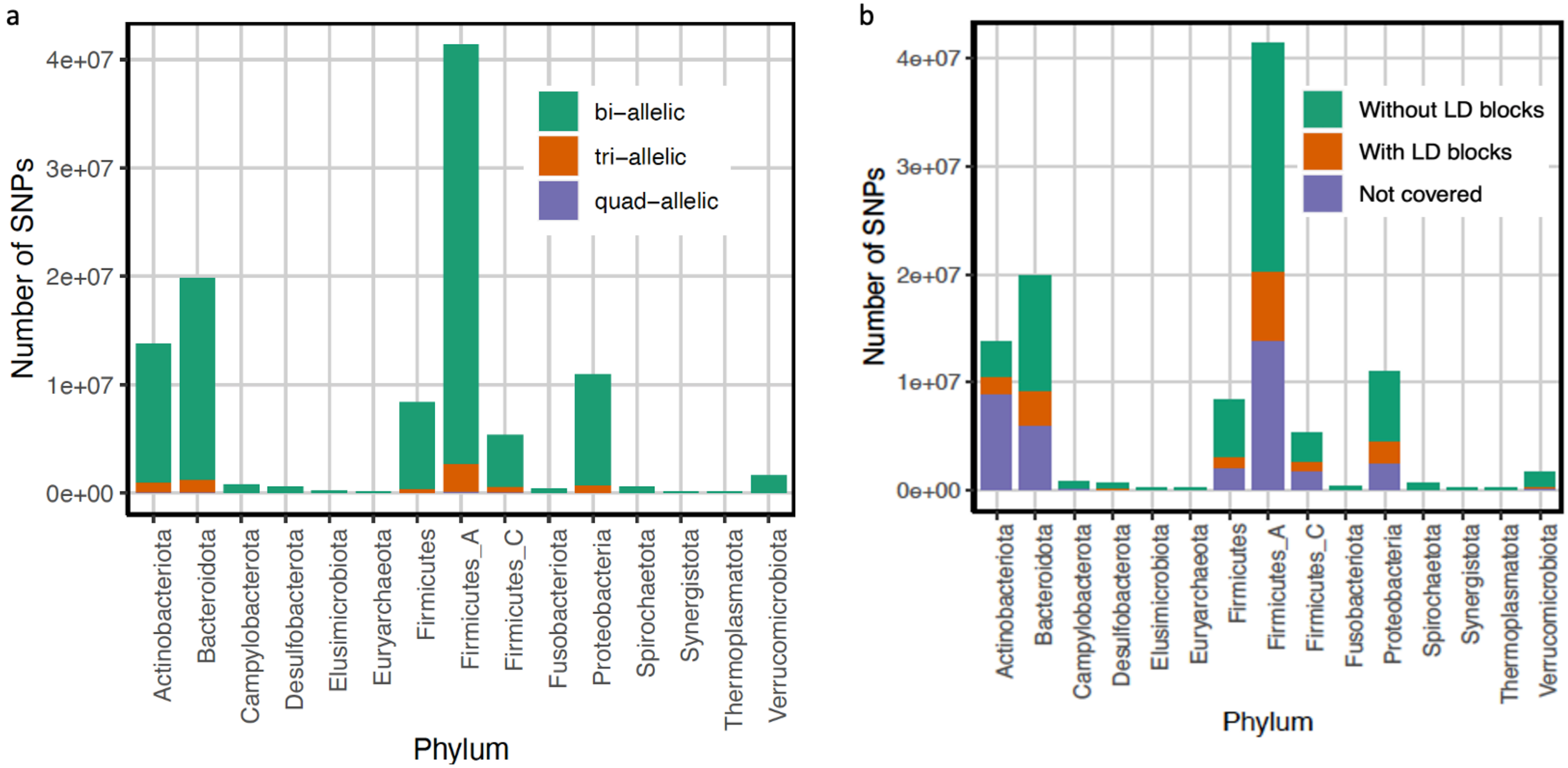
a
Number of SNPs
4e+07
3e+07
2e+07
1e+07
0e+00
bi–allelic
tri–allelic
quad–allelic
Actinobacteriota
Bacteroidota
Campylobacterota
Desulfobacterota
Elusimicrobiota
Euryarchaeota
Firmicutes
Firmicutes_A
Firmicutes_C
Fusobacteriota
Proteobacteria
Spirochaetota
Synergistota
Thermoplasmatota
Verrucomicrobiota
Phylum
b
Number of SNPs
4e+07
3e+07
2e+07
1e+07
0e+00
Without LD blocks
With LD blocks
Not covered
Actinobacteriota
Bacteroidota
Campylobacterota
Desulfobacterota
Elusimicrobiota
Euryarchaeota
Firmicutes
Firmicutes_A
Firmicutes_C
Fusobacteriota
Proteobacteria
Spirochaetota
Synergistota
Thermoplasmatota
Verrucomicrobiota
Phylum

**Figure S4**

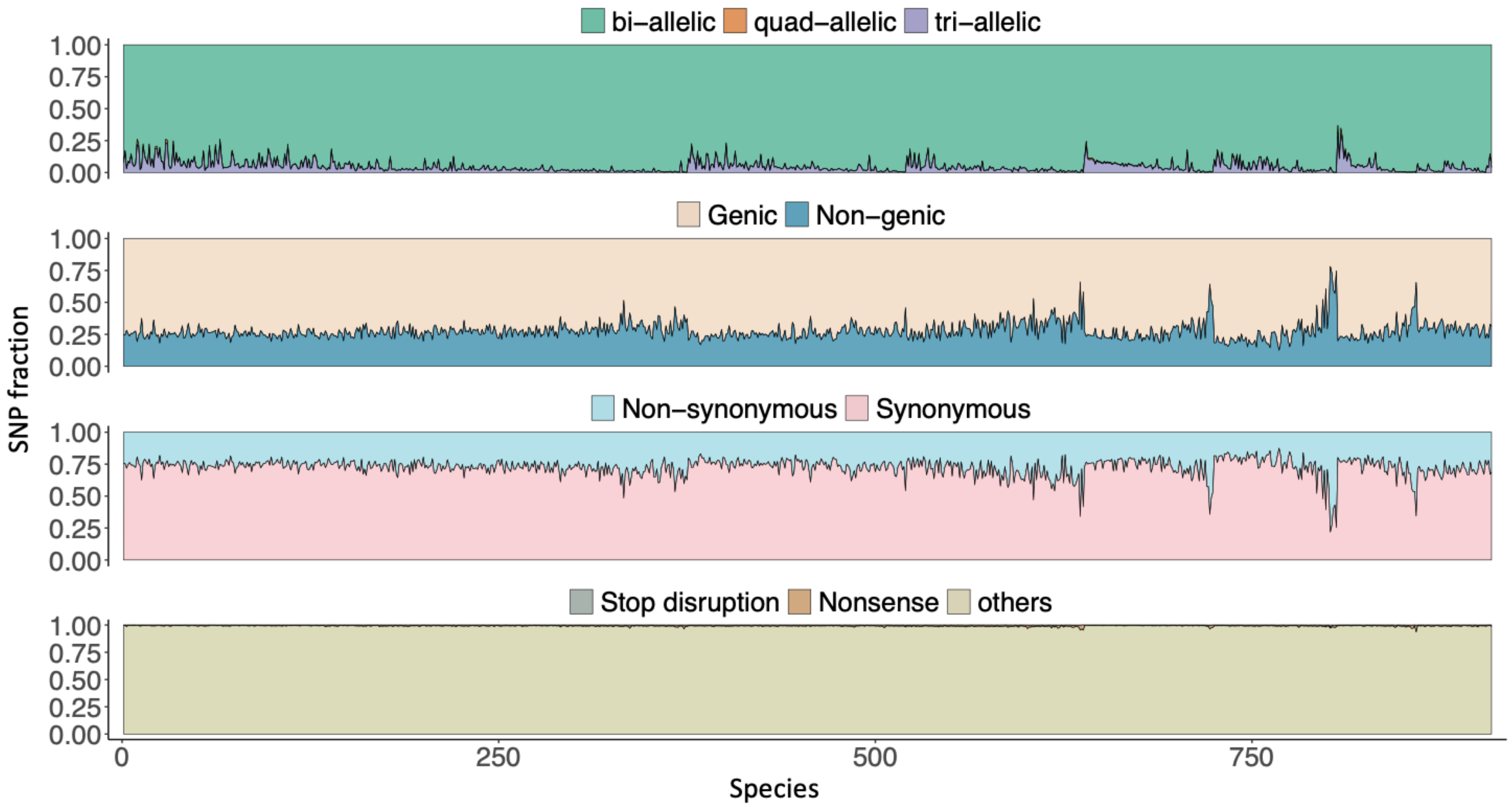
bi−allelic
quad−allelic
tri−allelic
Genic
Non−genic
Non−synonymous
Synonymous
Stop disruption
Nonsense
others
SNP fraction
1.00
0.75
0.50
0.25
0.00
0
250
500
750
Species

**Figure S5**

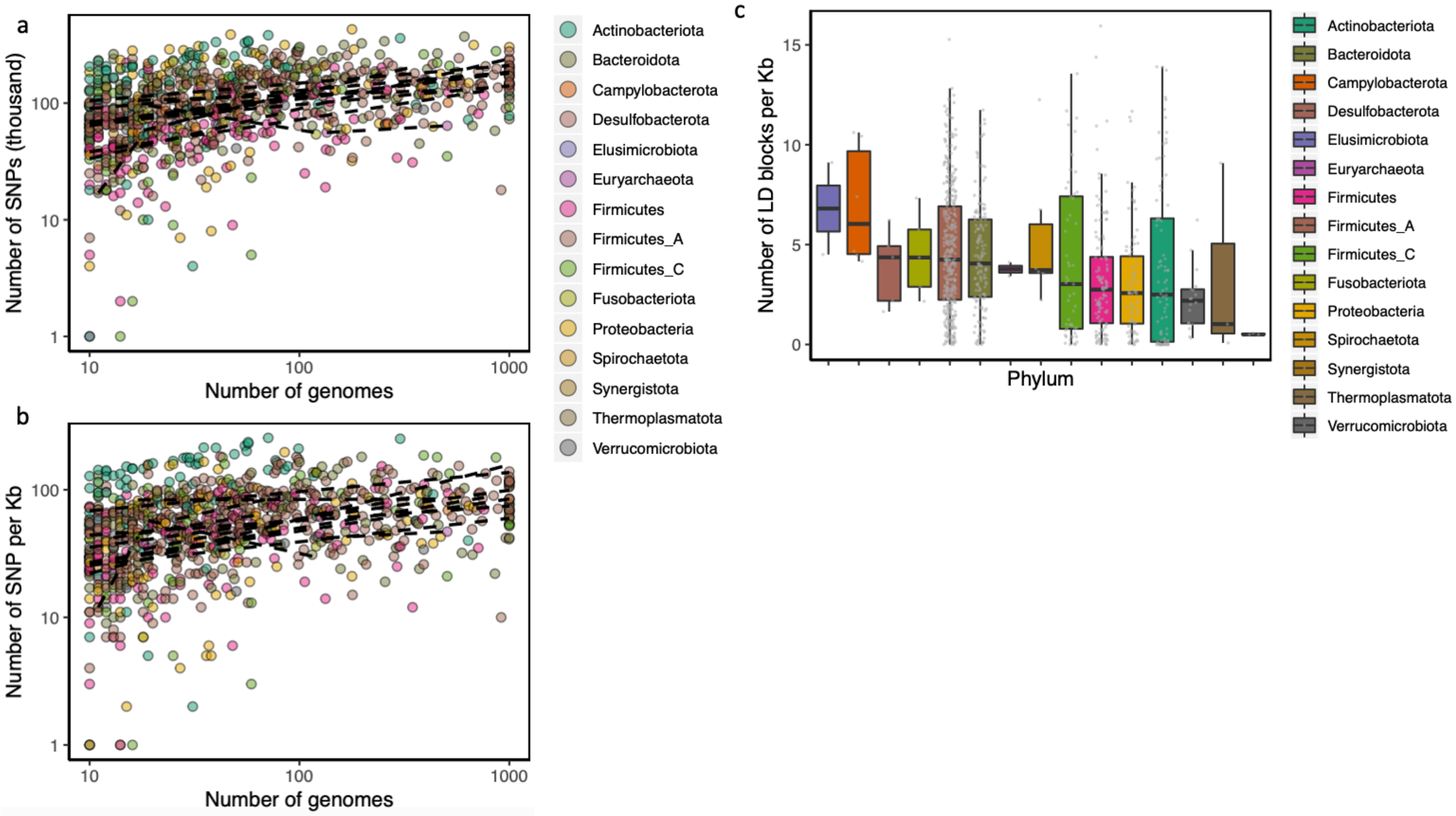
a
Number of SNPs (thousand)
Number of genomes
b
Number of SNP per Kb
Number of genomes
Actinobacteriota
Bacteroidota
Campylobacterota
Desulfobacterota
Elusimicrobiota
Euryarchaeota
Firmicutes
Firmicutes_A
Firmicutes_C
Fusobacteriota
Proteobacteria
Spirochaetota
Synergistota
Thermoplasmatota
Verrucomicrobiota
c
Number of LD blocks per Kb
Phylum
Actinobacteriota
Bacteroidota
Campylobacterota
Desulfobacterota
Elusimicrobiota
Euryarchaeota
Firmicutes
Firmicutes_A
Firmicutes_C
Fusobacteriota
Proteobacteria
Spirochaetota
Synergistota
Thermoplasmatota
Verrucomicrobiota

**Figure S6**

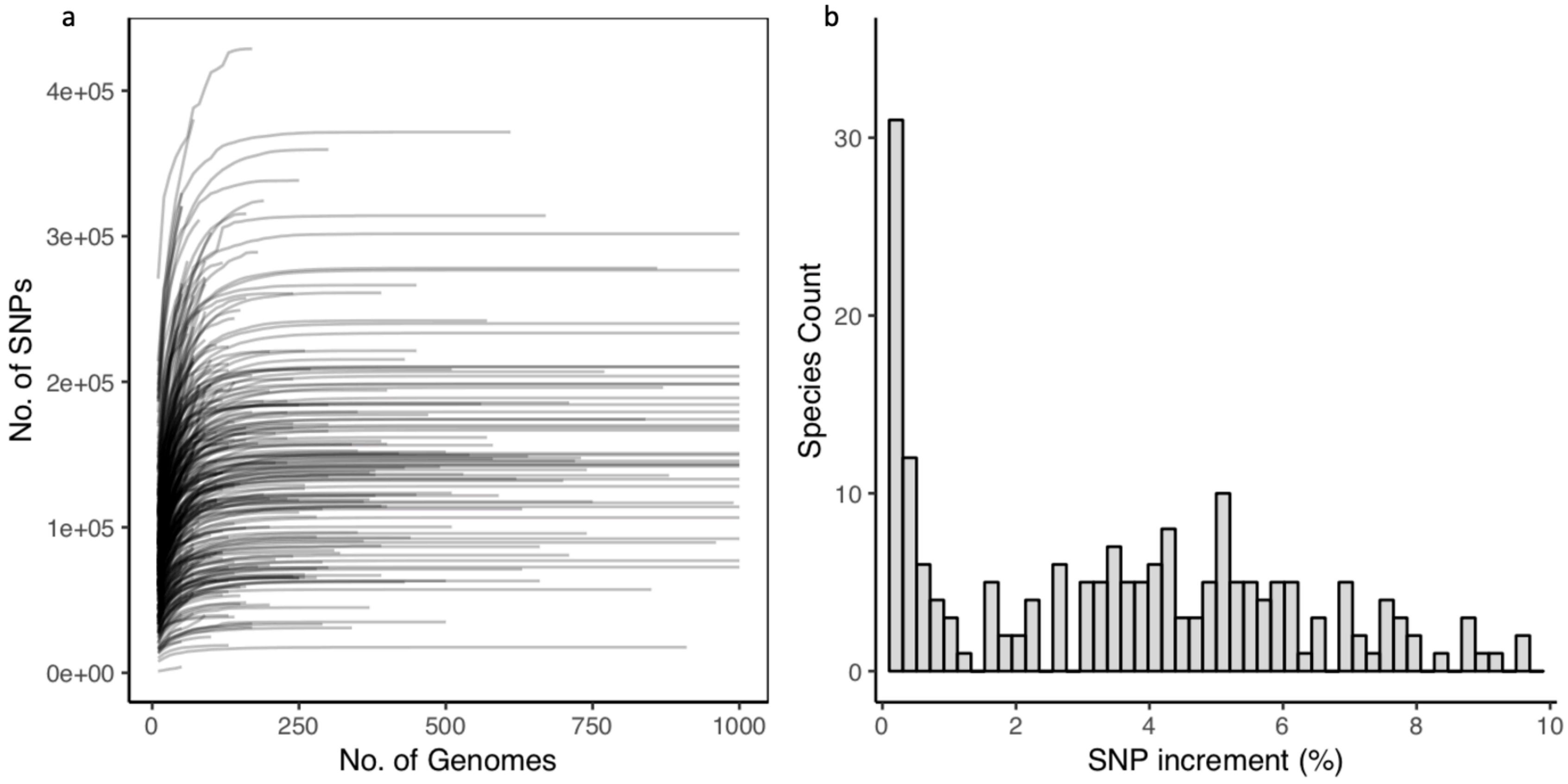
a
No. of SNPs
4e+05
3e+05
2e+05
1e+05
0e+00
0
250
500
750
1000
No. of Genomes
b
Species Count
30
20
10
0
0
2
4
6
8
10
SNP increment (%)

**Figure S7**

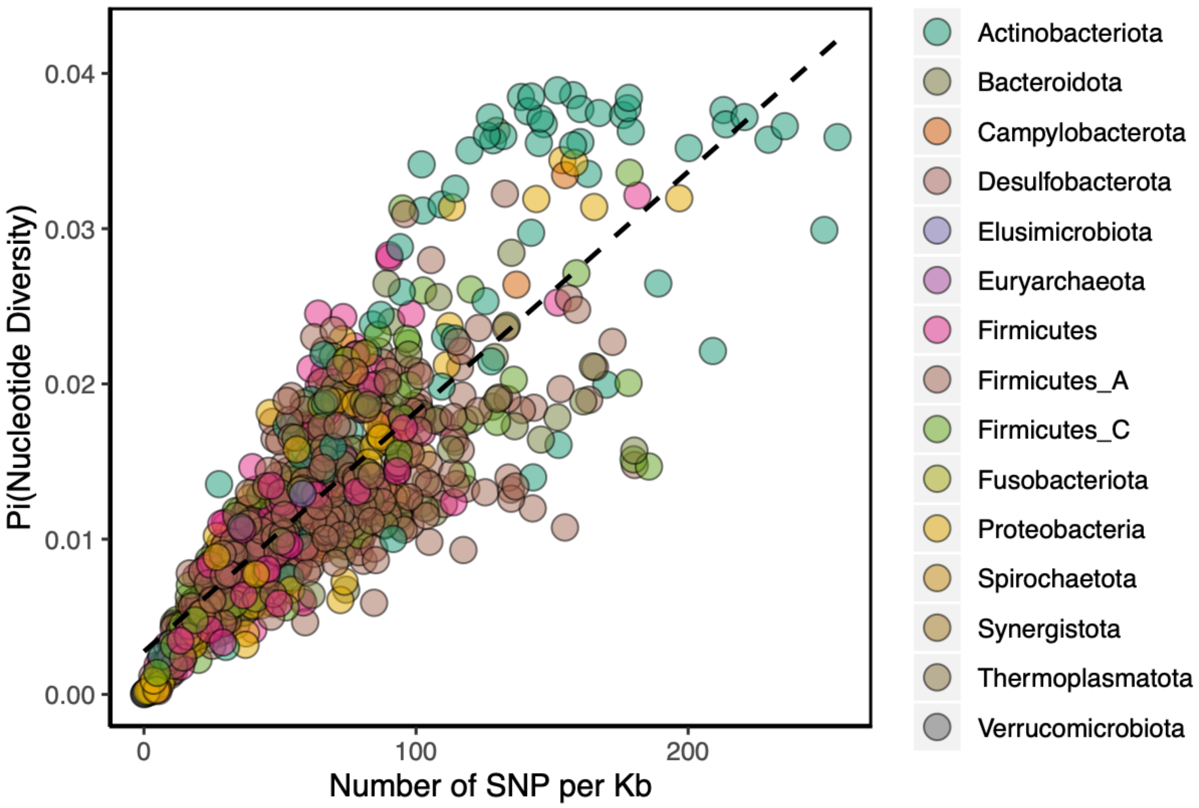
0.04
0.03
0.02
0.01
0.00
Pi(Nucleotide Diversity)
0
100
200
Number of SNP per Kb
Actinobacteriota
Bacteroidota
Campylobacterota
Desulfobacterota
Elusimicrobiota
Euryarchaeota
Firmicutes
Firmicutes_A
Firmicutes_C
Fusobacteriota
Proteobacteria
Spirochaetota
Synergistota
Thermoplasmatota
Verrucomicrobiota

**Figure S8**

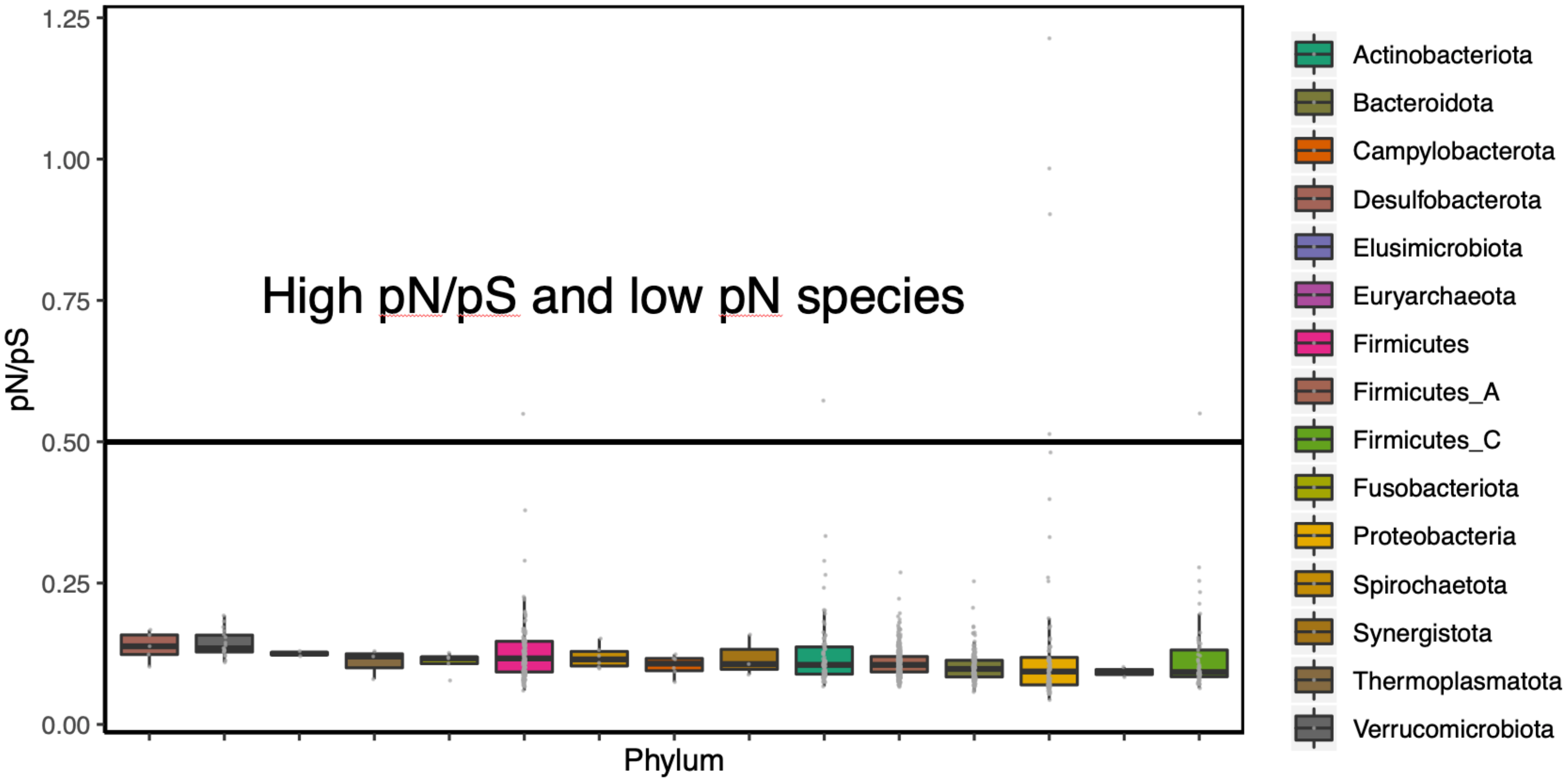

1.25
1.00
0.75
0.50
0.25
0.00
pN/pS
High pN/pS and low pN species
Phylum
Actinobacteriota
Bacteroidota
Campylobacterota
Desulfobacterota
Elusimicrobiota
Euryarchaeota
Firmicutes
Firmicutes_A
Firmicutes_C
Fusobacteriota
Proteobacteria
Spirochaetota
Synergistota
Thermoplasmatota
Verrucomicrobiota

**Figure S9**

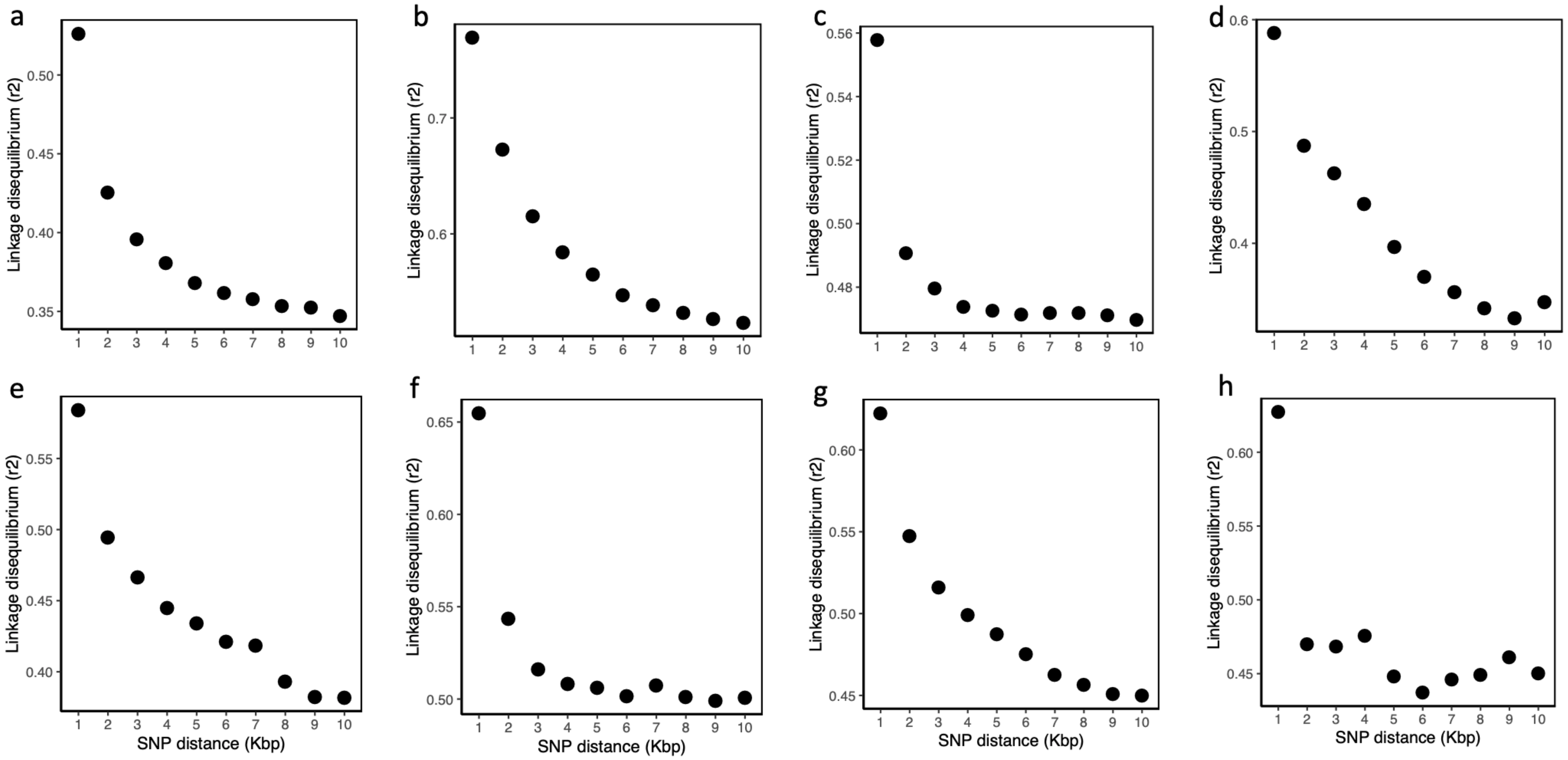

**Figure S10**

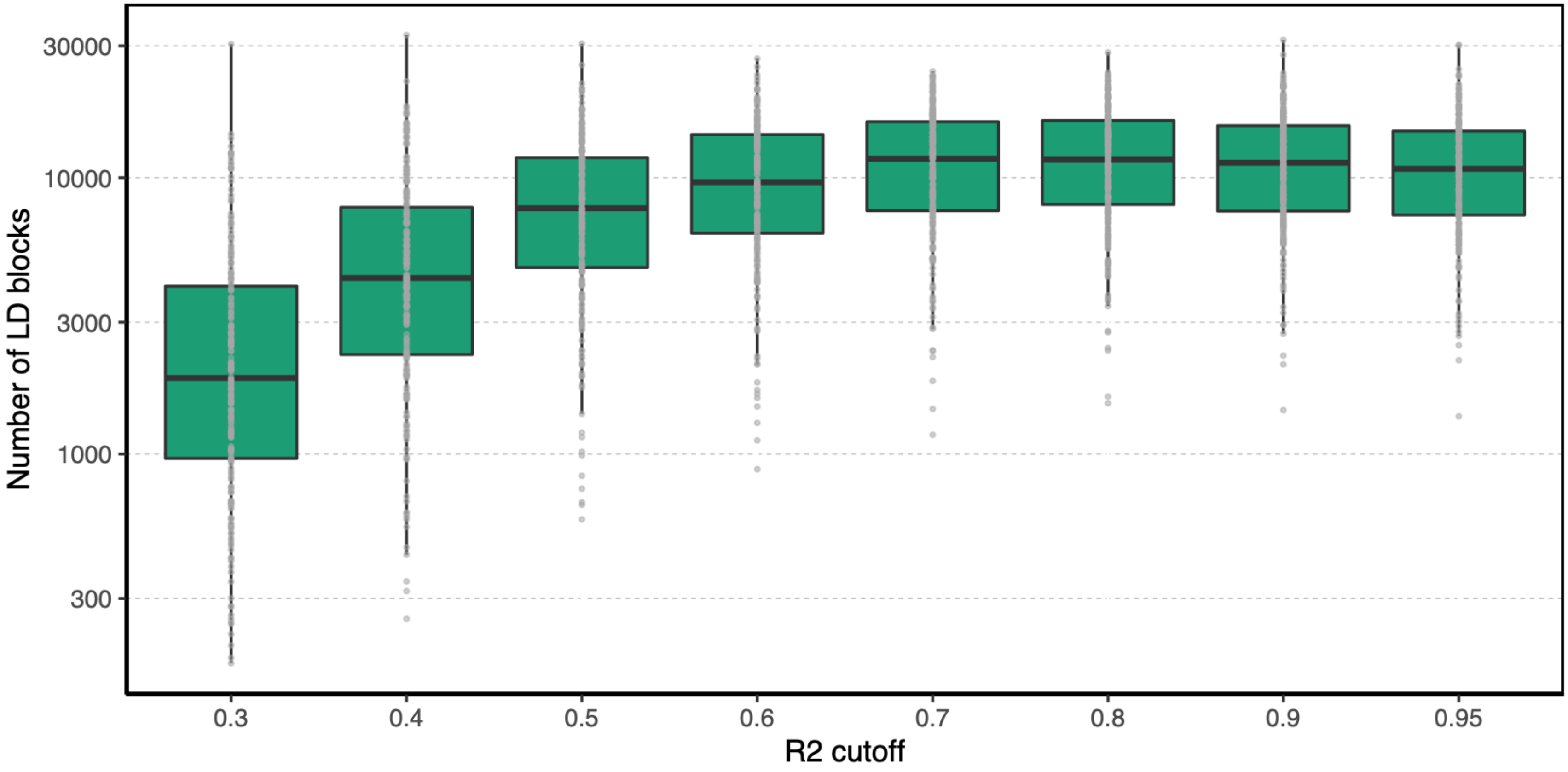

30000
10000
3000
1000
300
Number of LD blocks
0.3
0.4
0.5
0.6
0.7
0.8
0.9
0.95
R2 cutoff

**Figure S11**

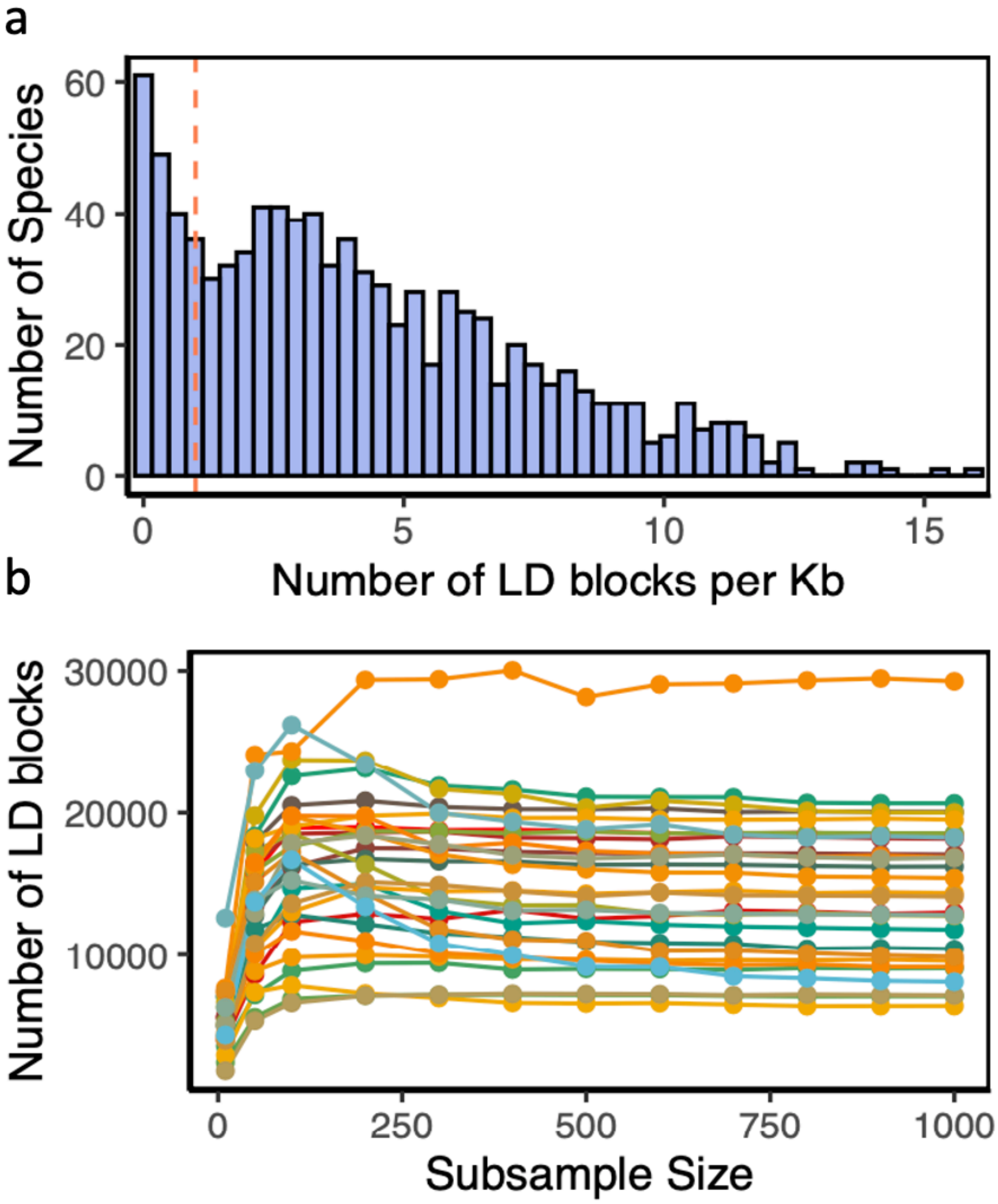
a
Number of Species
60
40
20
0
0
5
10
15
Number of LD blocks per Kb
b
Number of LD blocks
30000
20000
10000
0
250
500
750
1000
Subsample Size

**Figure S12**

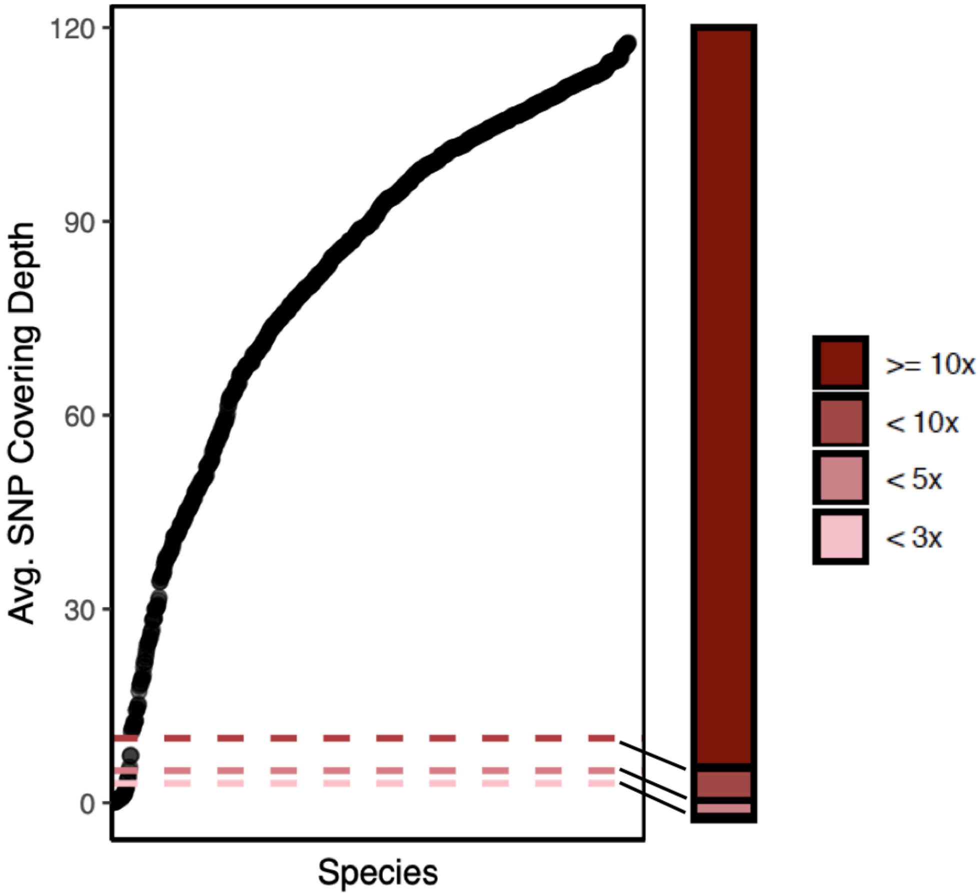

**Figure S13**

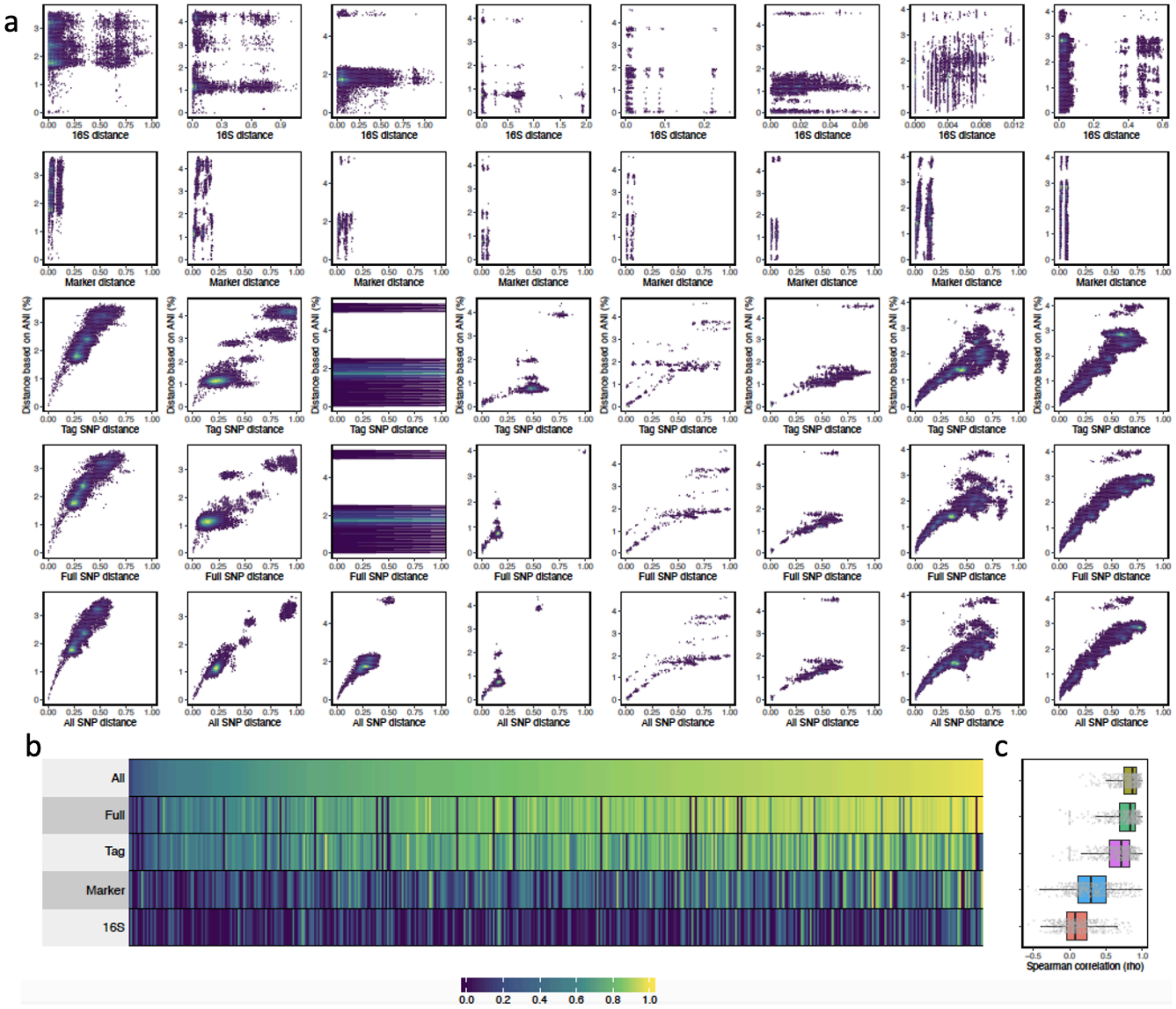
a
16S distance
Marker distance
Tag SNP distance
Full SNP distance
All SNP distance
Distance based on ANI (%)
b
All
Full
Tag
Marker
16S
c
Spearman correlation (rho)
0.0
0.2
0.4
0.6
0.8
1.0

### Figure S14

a

| SNP center | SNP offset |
| --- | --- |
| ..............................GGTCTAGGCGCAATGTAACGCTTTTATCGCT | 0 |
| ..........................TCTTGGTCTAGGCGCAATGTAACGCTTTTAT.... | 4 |
| ....................GCTTAGTCTTGGTCTAGGCGCAATGTAACGC.......... | 10 |
| ...............TTAGCGCTTAGTCTTGGTCTAGGCGCAATGT............... | 15 |
| ............GACTTAGCGCTTAGTCTTGGTCTAGGCGCAA.................. | 18 |
| .........CTAGACTTAGCGCTTAGTCTTGGTCTAGGCG..................... | 21 |
| CAACTATTCCTAGACTTAGCGCTTAGTCTTGGTCTAGGCGCAATGTAACGCTTTTATCGCT | Major sc-span |
| CAACTATTCCTAGACTTAGCGCTTAGTCTTCGTCTAGGCGCAATGTAACGCTTTTATCGCT | Minor sc-span |
| ....................GCTTAGTCTTCGTCTAGGCGCAATGTAACGC.......... | 10 |
| ..............CTTAGCGCTTAGTCTTCGTCTAGGCGCAATG................ | 16 |
| .........CTAGACTTAGCGCTTAGTCTTCGTCTAGGCG..................... | 21 |
| .....ATTCCTAGACTTAGCGCTTAGTCTTCGTCTA......................... | 25 |
| ....TATTCCTAGACTTAGCGCTTAGTCTTCGTCT.......................... | 26 |

b

| SNP center | SNP offset |
| --- | --- |
| ..........................TCTTGGTCTAGGCGCAATGTAACGCTTTTAT.... | 4 |
| ....................GCTTAGTCTTGGTCTAGGCGCAATGTAACGC.......... | 10 |
| ...............TTAGCGCTTAGTCTTGGTCTAGGCGCAATGT............... | 15 |
| ............GACTTAGCGCTTAGTCTTGGTCTAGGCGCAA.................. | 18 |
| .........CTAGACTTAGCGCTTAGTCTTGGTCTAGGCG..................... | 21 |
| CAACTATTCCTAGACTTAGCGCTTAGTCTTGGTCTAGGCGCAATGTAACGCTTTTATCGCT | Major sc-span |
| CAACTATTCCTAGACTTAGCGCTTAGTCTTCGTCTAGGCGCAATGTAACGCTTTTATCGCT | Minor sc-span |
| ....................GCTTAGTCTTCGTCTAGGCGCAATGTAACGC.......... | 10 |
| ..............CTTAGCGCTTAGTCTTCGTCTAGGCGCAATG................ | 16 |
| CAACTATTCCTAGACTTAGCTCTTAGTCTTCGTCTAGGCGCAATGTAACGCTTTTATCGCT | Minor sc-span′ |
| .........CTAGACTTAGCTCTTAGTCTTCGTCTAGGCG..................... | 21 |
| .....ATTCCTAGACTTAGCTCTTAGTCTTCGTCTA......................... | 25 |
| ....TATTCCTAGACTTAGCTCTTAGTCTTCGTCT.......................... | 26 |

**Figure S15**

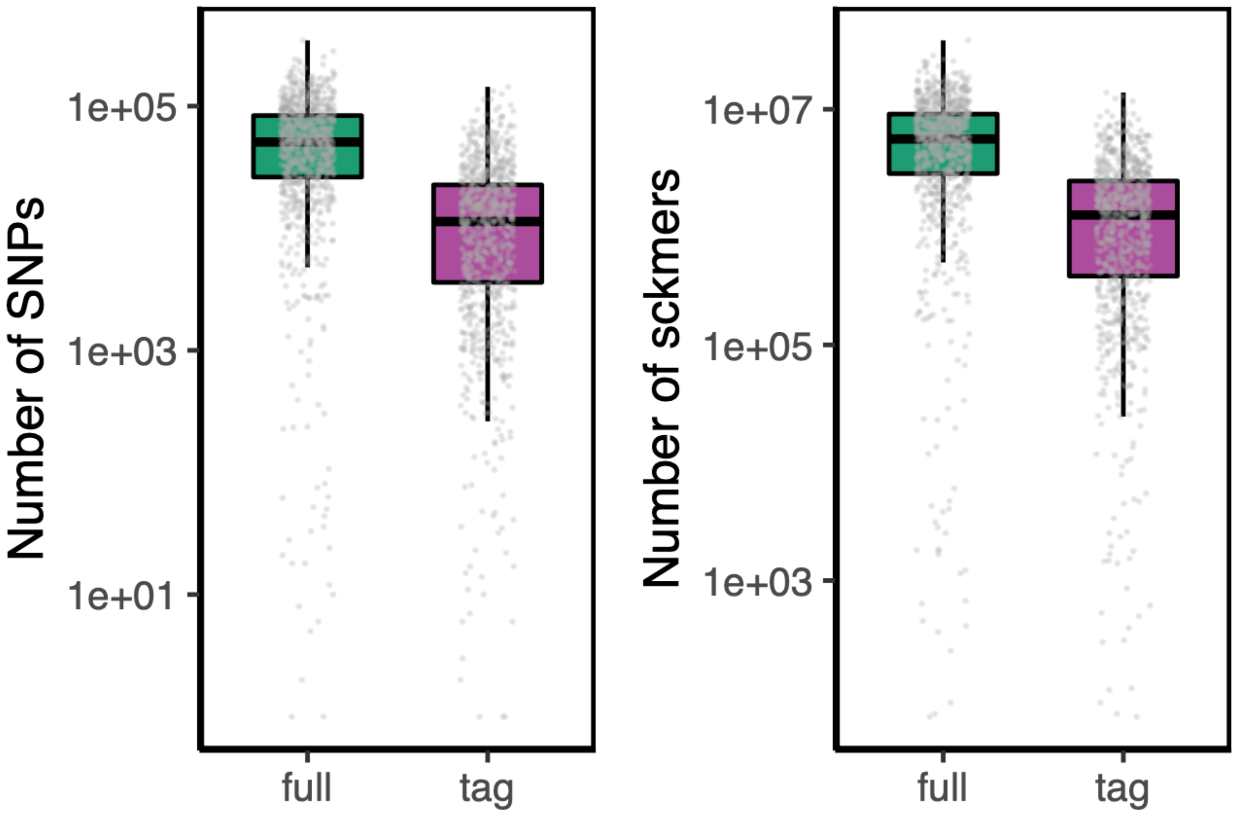

Number of SNPs
1e+05
1e+03
1e+01
full
tag
Number of sckmers
1e+07
1e+05
1e+03
full
tag

### Figure S16

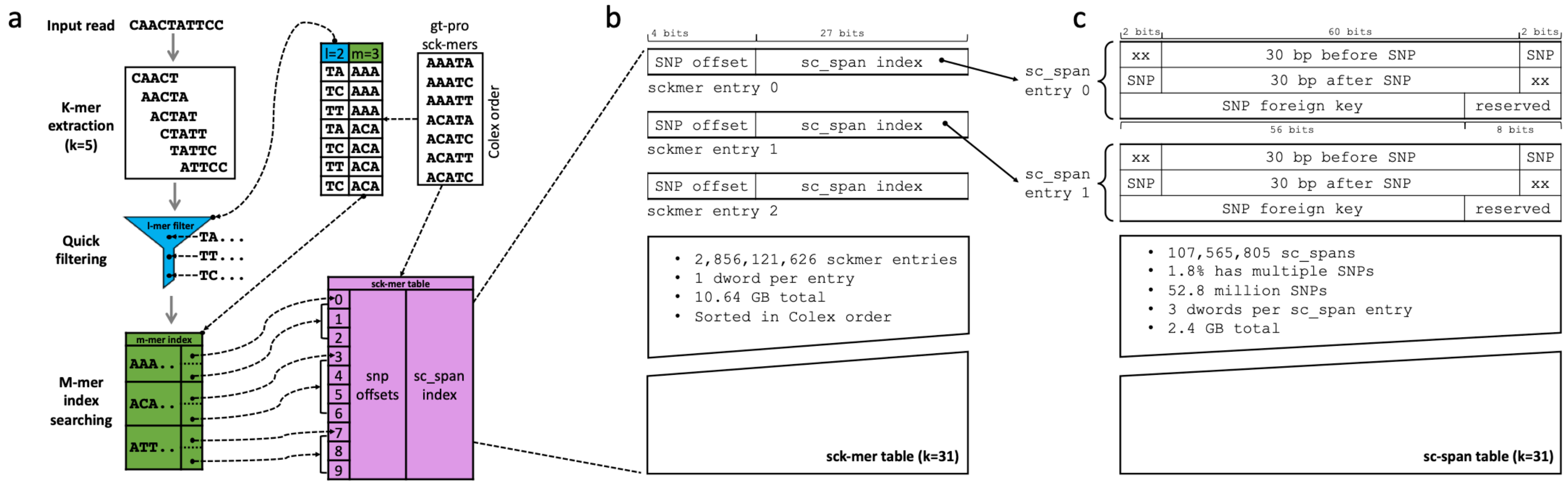

a
Input read
CAACTATTCC
K-mer extraction (k=5)
CAACT
AACTA
ACTAT
CTATT
TATTC
ATTCC
Quick filtering
l-mer filter
TA...
TT...
TC...
M-mer index searching
m-mer index
AAA..
ACA..
ATT..
l=2
m=3
gt-pro sck-mers
AAATA
AAATC
AAATT
ACATA
ACATC
ACATT
ACATC
Colex order
sck-mer table
snp offsets
sc_span index
b
4 bits
27 bits
SNP offset
sc_span index
sckmer entry 0
sckmer entry 1
sckmer entry 2
2,856,121,626 sckmer entries
1 dword per entry
10.64 GB total
Sorted in Colex order
sck-mer table (k=31)
c
2 bits
60 bits
2 bits
sc_span entry 0
sc_span entry 1
xx
30 bp before SNP
SNP
30 bp after SNP
SNP foreign key
reserved
56 bits
8 bits
107,565,805 sc_spans
1.8% has multiple SNPs
52.8 million SNPs
3 dwords per sc_span entry
2.4 GB total
sc-span table (k=31)

**Figure S17**

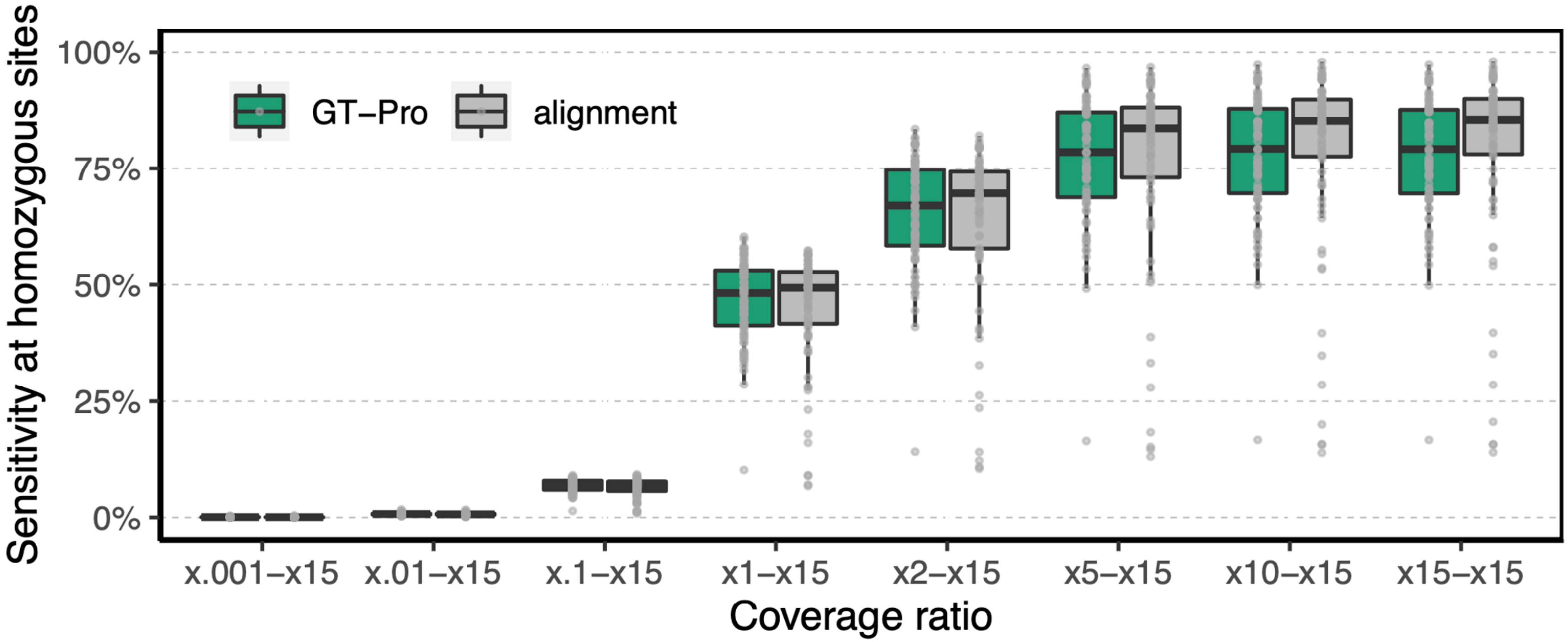

Sensitivity at homozygous sites
100%
75%
50%
25%
0%
GT−Pro
alignment
x.001−x15
x.01−x15
x.1−x15
x1−x15
x2−x15
x5−x15
x10−x15
x15−x15
Coverage ratio

**Figure S18**

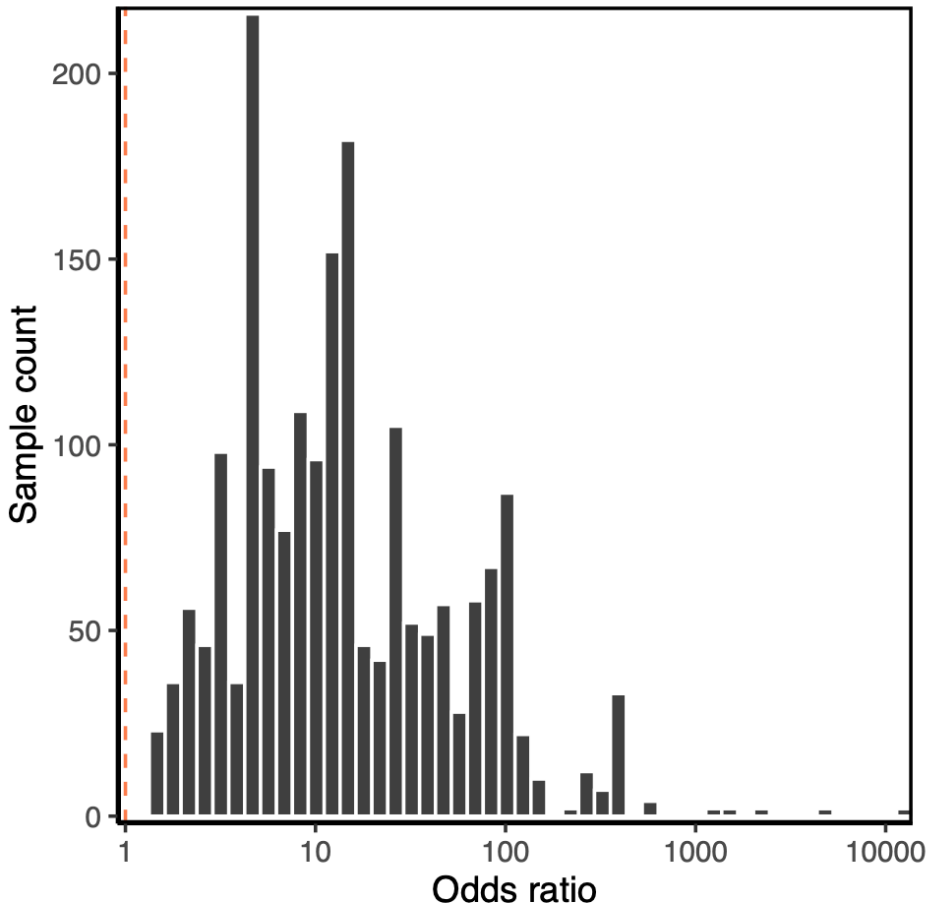

**Figure S19**

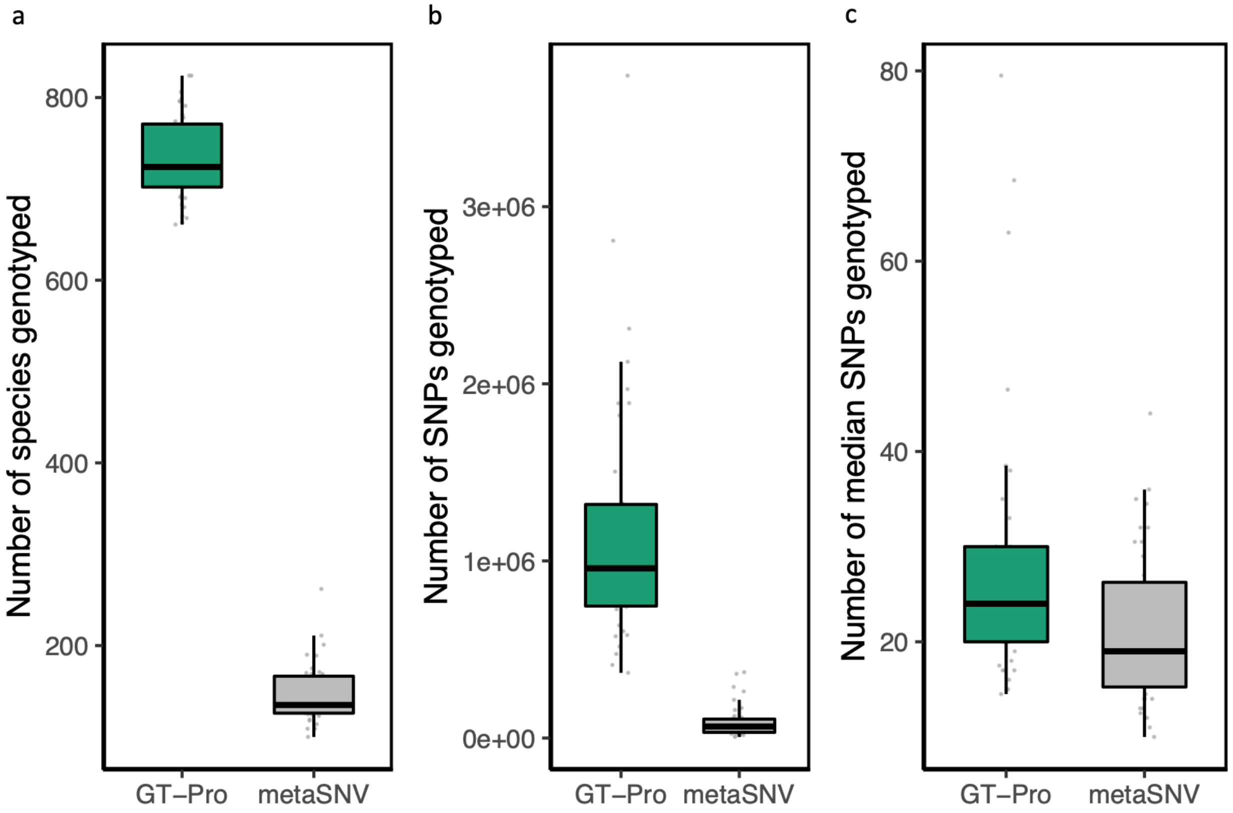

a
b
c
Number of species genotyped
800
600
400
200
GT−Pro
metaSNV
Number of SNPs genotyped
3e+06
2e+06
1e+06
0e+00
GT−Pro
metaSNV
Number of median SNPs genotyped
80
60
40
20
GT−Pro
metaSNV

**Figure S20**

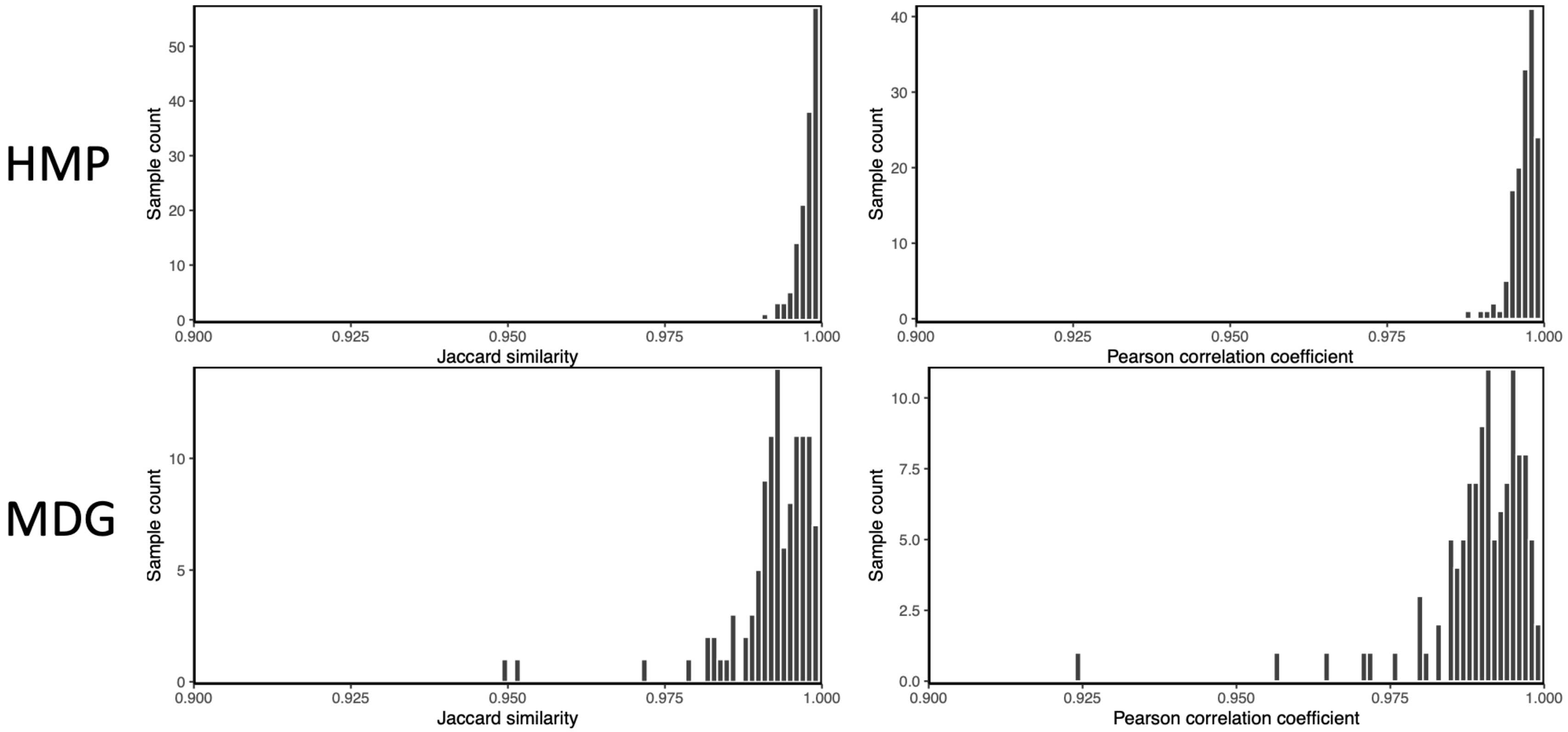
HMP
MDG
Sample count
0
10
20
30
40
50
0.900
0.925
0.950
0.975
1.000
Jaccard similarity
Sample count
0
10
20
30
40
0.900
0.925
0.950
0.975
1.000
Pearson correlation coefficient
Sample count
0
5
10
0.900
0.925
0.950
0.975
1.000
Jaccard similarity
Sample count
0.0
2.5
5.0
7.5
10.0
0.900
0.925
0.950
0.975
1.000
Pearson correlation coefficient

**Figure S21**

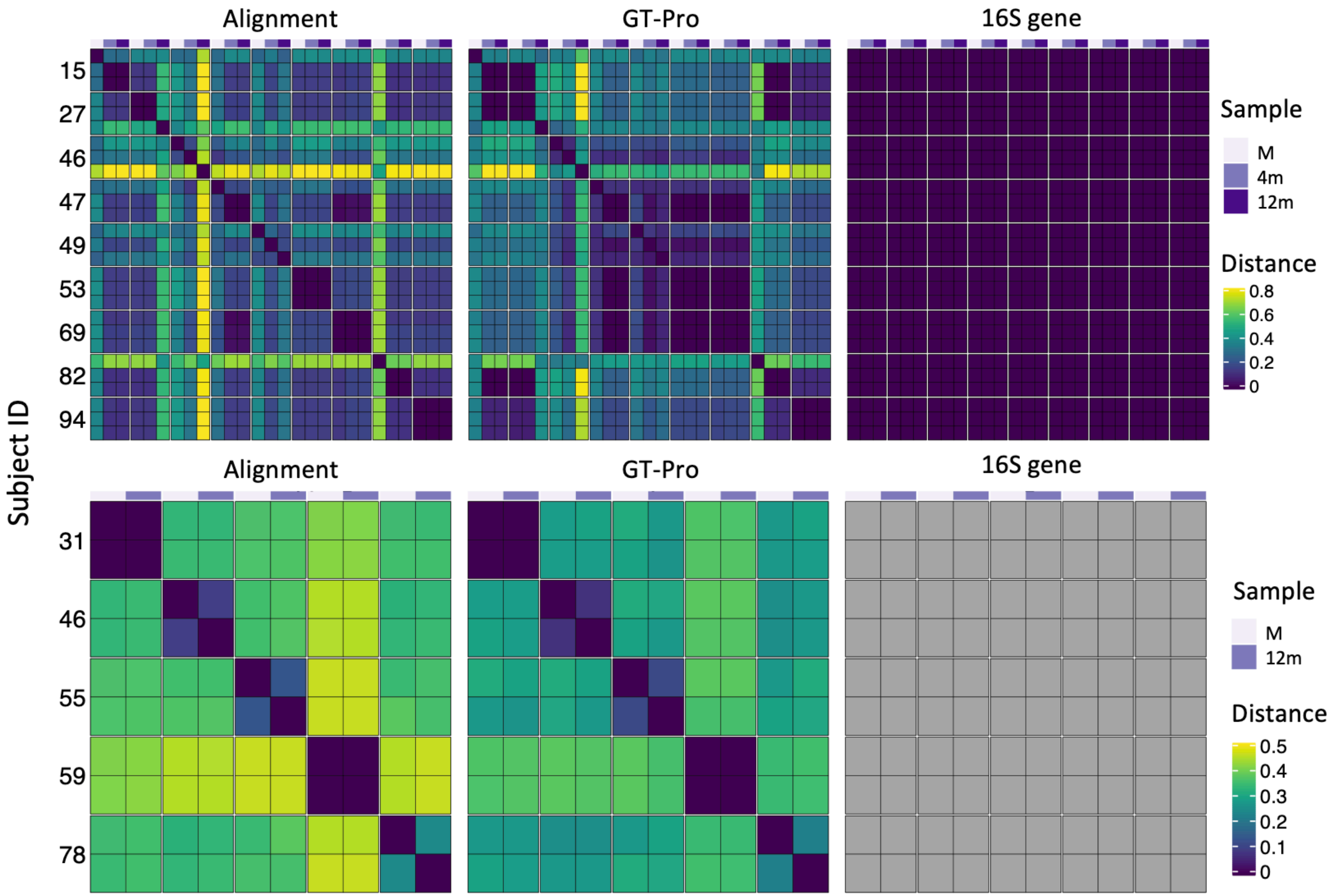

Alignment
GT-Pro
16S gene
Subject ID
15
27
46
47
49
53
69
82
94
Sample
M
4m
12m
Distance
0.8
0.6
0.4
0.2
0
Alignment
GT-Pro
16S gene
31
46
55
59
78
Sample
M
12m
Distance
0.5
0.4
0.3
0.2
0.1
0

### Figure S22

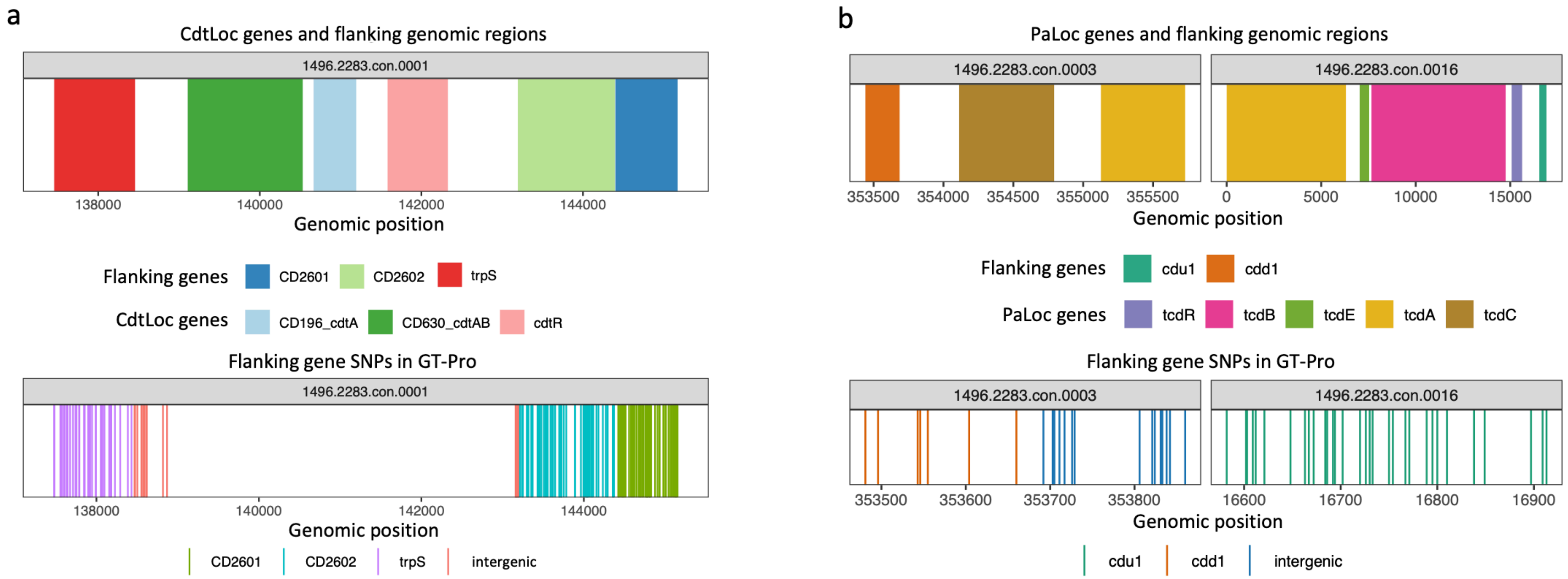
a
CdtLoc genes and flanking genomic regions
1496.2283.con.0001
138000
140000
142000
144000
Genomic position
Flanking genes
CD2601
CD2602
trpS
CdtLoc genes
CD196_cdtA
CD630_cdtAB
cdtR
Flanking gene SNPs in GT-Pro
1496.2283.con.0001
138000
140000
142000
144000
Genomic position
CD2601
CD2602
trpS
intergenic
b
PaLoc genes and flanking genomic regions
1496.2283.con.0003
1496.2283.con.0016
353500
354000
354500
355000
355500
0
5000
10000
15000
Genomic position
Flanking genes
cdu1
cdd1
PaLoc genes
tcdR
tcdB
tcdE
tcdA
tcdC
Flanking gene SNPs in GT-Pro
1496.2283.con.0003
1496.2283.con.0016
353500
353600
353700
353800
16600
16700
16800
16900
Genomic position
cdu1
cdd1
intergenic

**Figure S23**

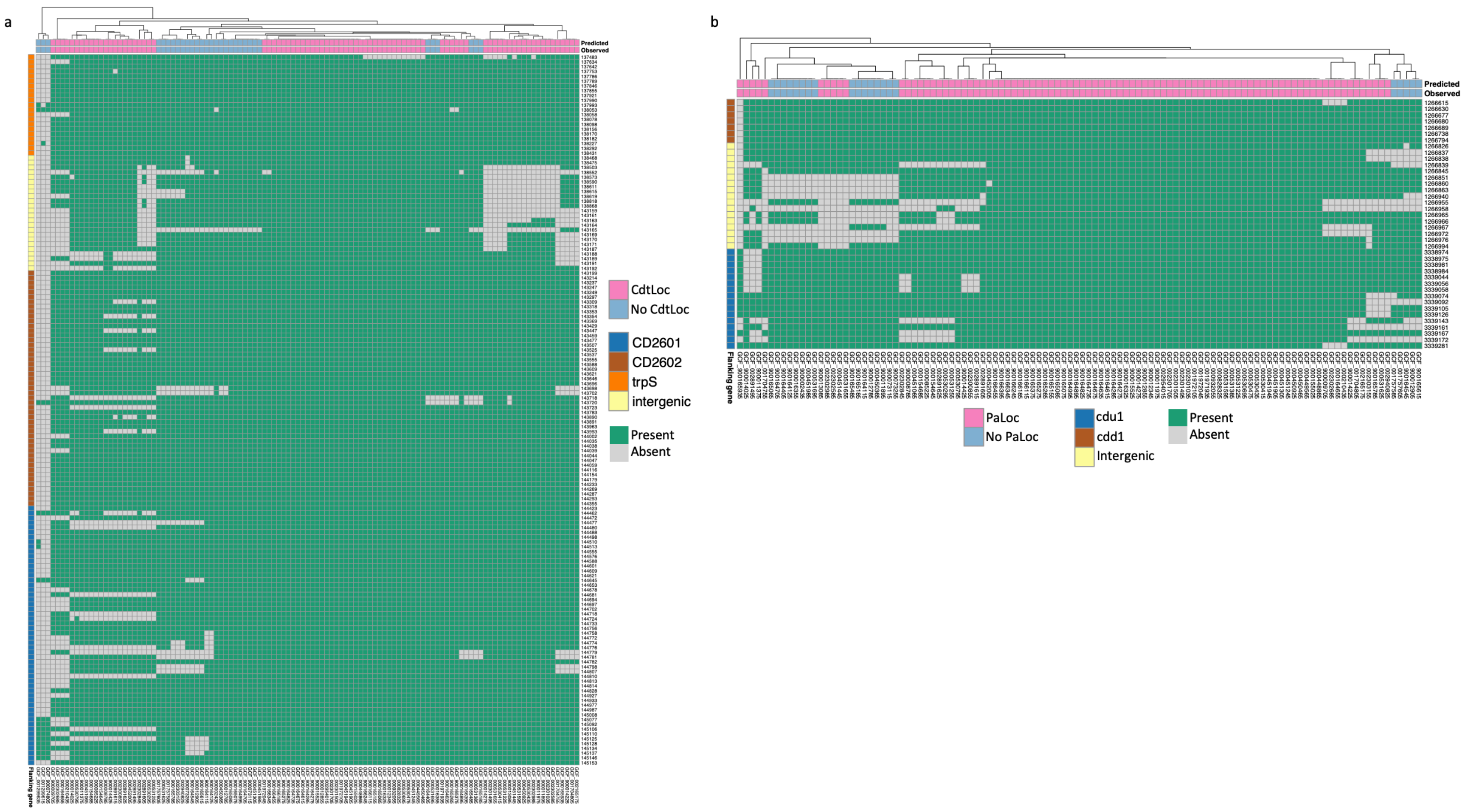
a
Predicted
Observed
Flanking gene
CdtLoc
No CdtLoc
CD2601
CD2602
trpS
intergenic
Present
Absent
b
Predicted
Observed
Flanking gene
PaLoc
No PaLoc
cdu1
cdd1
Intergenic
Present
Absent

**Figure S24**

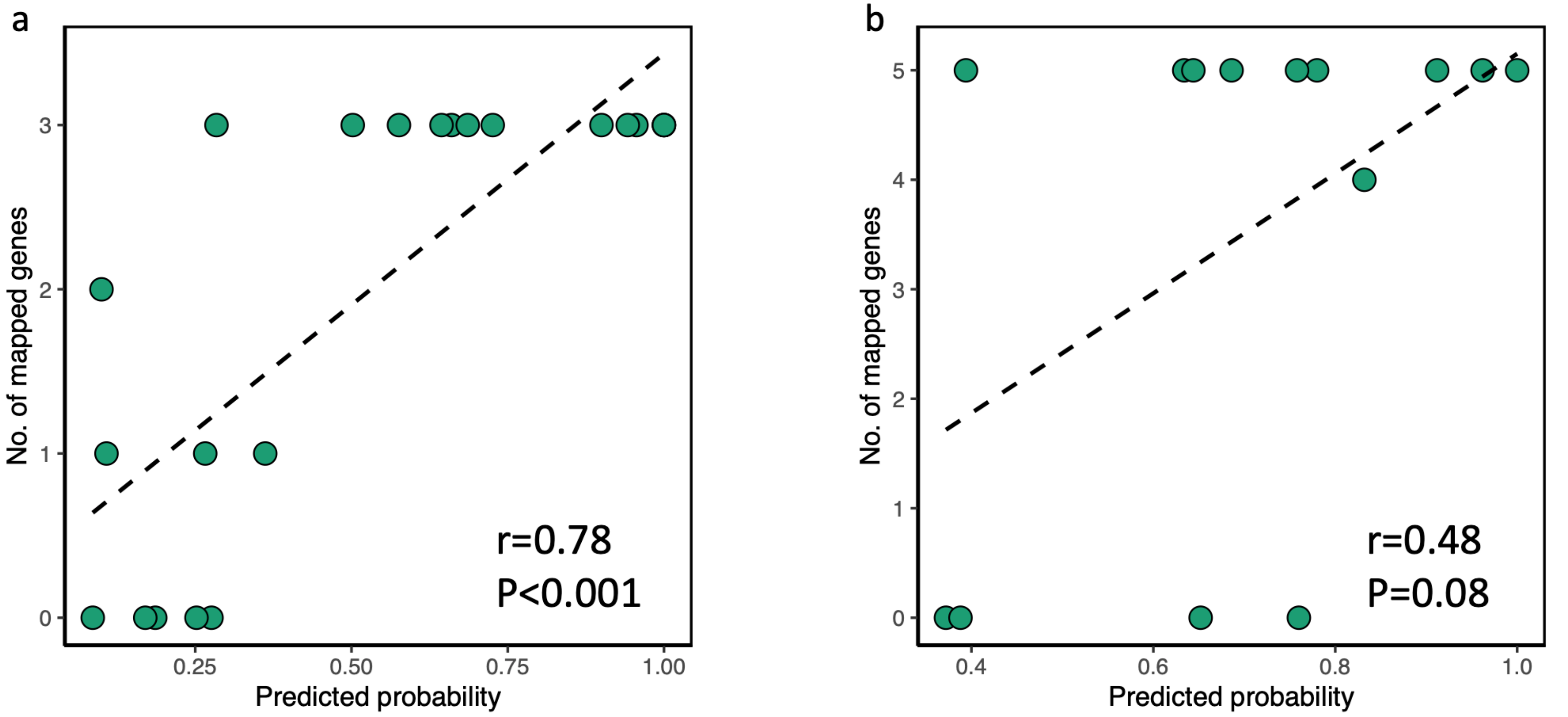
a
No. of mapped genes
Predicted probability
0.25
0.50
0.75
1.00
0
1
2
3
r=0.78
P<0.001
b
No. of mapped genes
Predicted probability
0.4
0.6
0.8
1.0
0
1
2
3
4
5
r=0.48
P=0.08

**Figure S25**

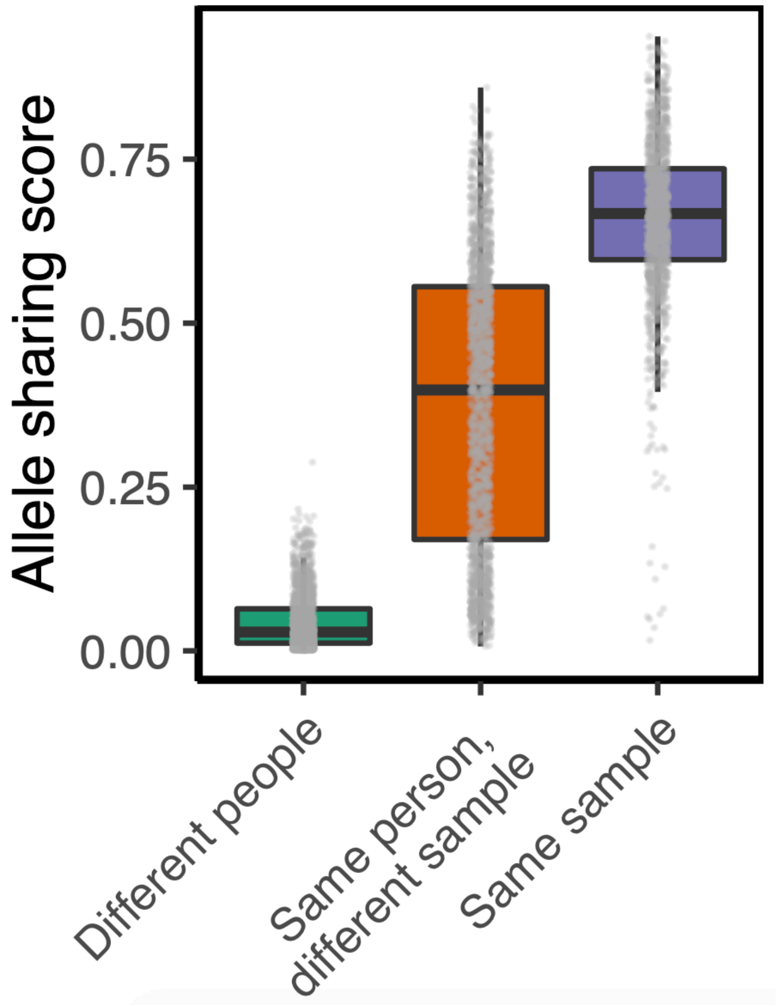

Allele sharing score
0.75
0.50
0.25
0.00
Different people
Same person, different sample
Same sample

**Figure S26**

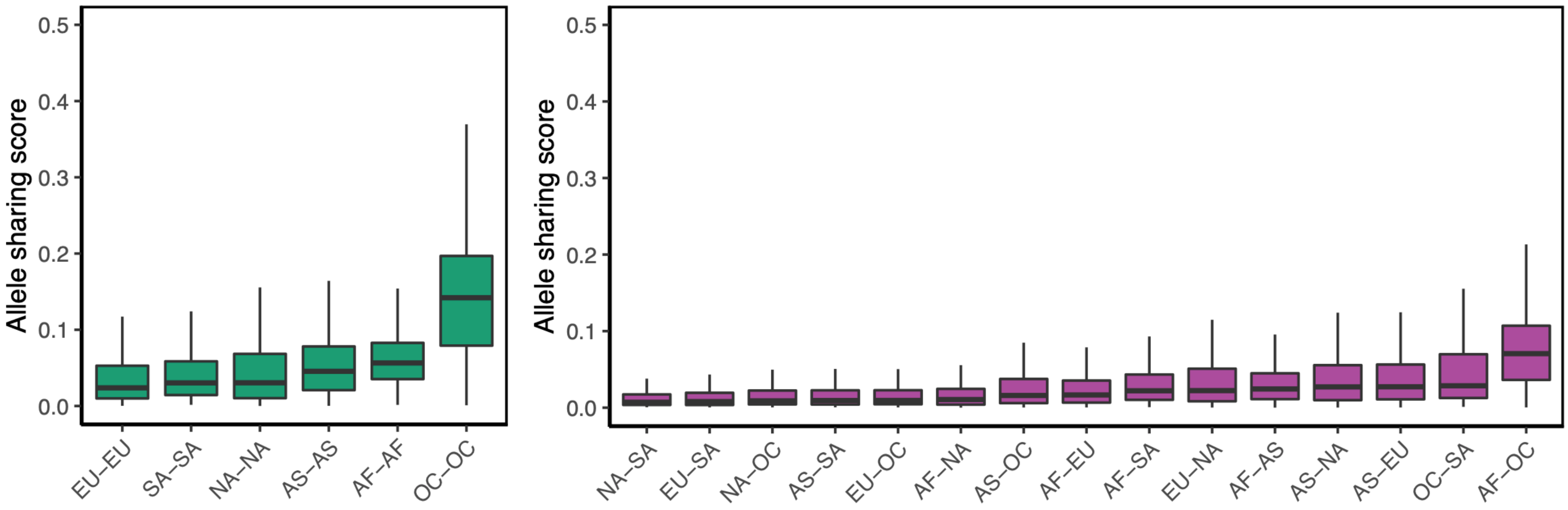
Allele sharing score
0.5
0.4
0.3
0.2
0.1
0.0
EU–EU
SA–SA
NA–NA
AS–AS
AF–AF
OC–OC
Allele sharing score
0.5
0.4
0.3
0.2
0.1
0.0
NA–SA
EU–SA
NA–OC
AS–SA
EU–OC
AF–NA
AS–OC
AF–EU
AF–SA
EU–NA
AF–AS
AS–NA
AS–EU
OC–SA
AF–OC

**Figure S27**

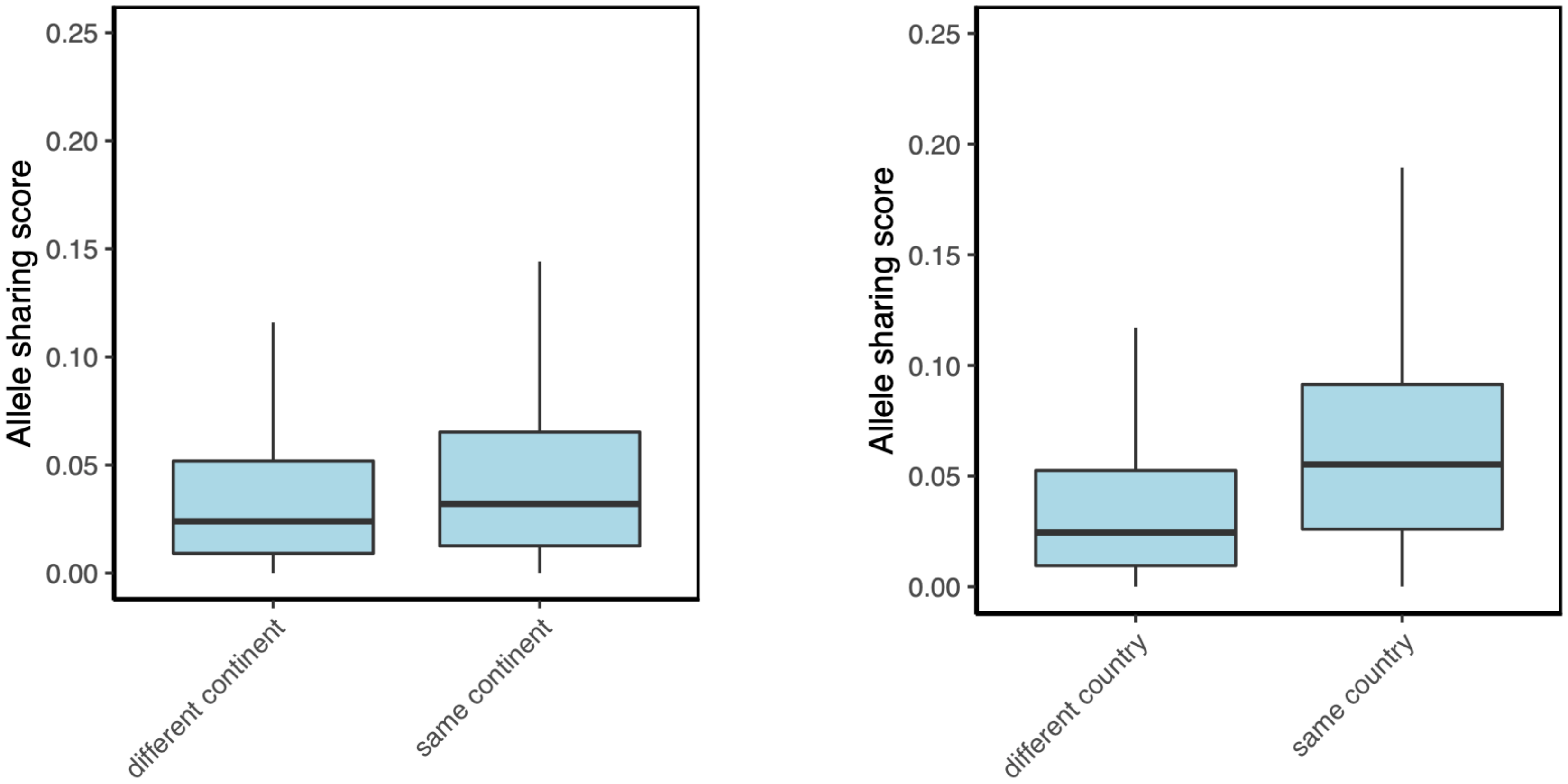

Allele sharing score
0.25
0.20
0.15
0.10
0.05
0.00
different continent
same continent
Allele sharing score
0.25
0.20
0.15
0.10
0.05
0.00
different country
same country

**Figure S28**

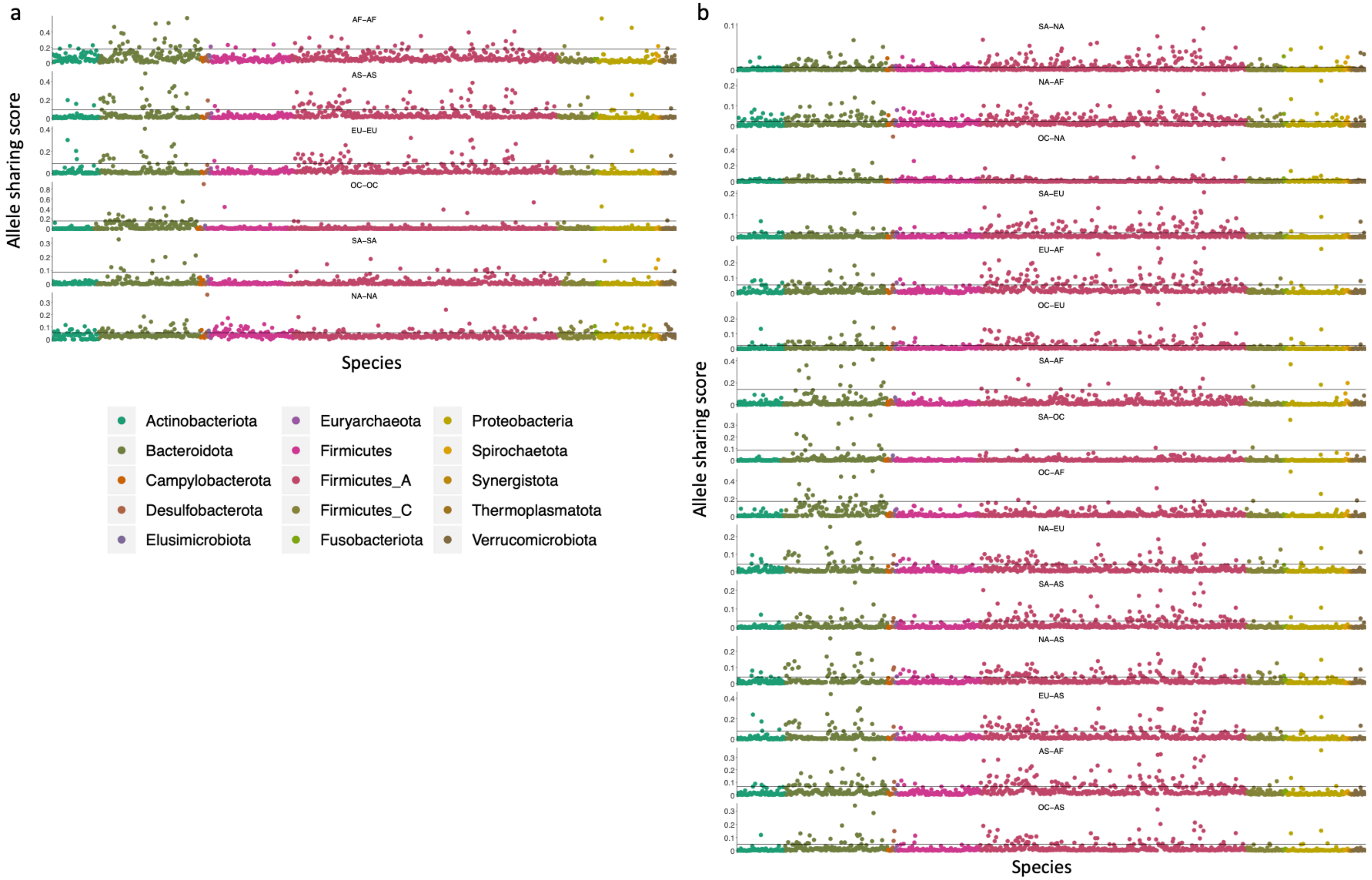
a
AF–AF
AS–AS
EU–EU
OC–OC
SA–SA
NA–NA
Allele sharing score
Species
Actinobacteriota
Bacteroidota
Campylobacterota
Desulfobacterota
Elusimicrobiota
Euryarchaeota
Firmicutes
Firmicutes_A
Firmicutes_C
Fusobacteriota
Proteobacteria
Spirochaetota
Synergistota
Thermoplasmatota
Verrucomicrobiota
b
SA–NA
NA–AF
OC–NA
SA–EU
EU–AF
OC–EU
SA–AF
SA–OC
OC–AF
NA–EU
SA–AS
NA–AS
EU–AS
AS–AF
OC–AS
Allele sharing score
Species
